# Psmd13, a proteosome regulatory subunit identified in miR-29a regulation during neurogenesis

**DOI:** 10.1101/2025.04.14.648858

**Authors:** Diji Kuriakose, Grant Morahan, Zhicheng Xiao

## Abstract

miR-29a is essential for neuronal development and implicated in neurodegenerative diseases, yet its regulatory mechanisms remain poorly understood. To identify upstream regulators of miR-29a expression, we leveraged the genetically diverse Collaborative Cross (CC) mouse strains and performed expression profiling and Quantitative Trait Loci (QTL) analysis, identifying a significant QTL on chromosome 7. Among ten candidate genes, Psmd13 emerged as a key regulator, with RNAi-mediated knockdown in mouse neural precursor cells (mNPCs) leading to enhanced neuronal differentiation and increased miR-29a expression in the undifferentiated state but decreased expression upon differentiation. Co-immunoprecipitation assays revealed that Psmd13 interacts with Dicer, modulating miR-29a levels in a differentiation-dependent manner. Chromatin immunoprecipitation sequencing (ChIP-seq) demonstrated Psmd13-Dicer co-binding at genomic loci, including miR-29a, suggesting a role in chromatin accessibility and transcriptional regulation. Proteasome inhibition using MG132 reduced Psmd13 and Dicer levels, downregulating miR-29a and impairing neuronal differentiation. These findings indicate that differentiation dynamically alters miR-29a transcription through Psmd13-Dicer interactions, positioning Psmd13 as a critical mediator of miR regulation and neurodevelopmental homeostasis.

## Introduction

MicroRNAs (miRs) are small, non-coding RNA molecules, typically 20-24 nucleotides in length, that play a pivotal role in gene regulation by targeting messenger RNAs (mRNAs). They primarily bind to the 3’ untranslated regions (UTRs) of mRNAs, leading to either the inhibition of translation or the degradation of the mRNA^1^. By modulating the expression of various protein-coding genes, miRs influence essential biological processes such as cell proliferation, differentiation, apoptosis, and stress responses. This intricate regulation is critical in maintaining cellular homeostasis, and disruptions in miR expression are often linked to diseases like cancer, cardiovascular disorders, and neurodegenerative conditions. Despite their importance, the mechanisms controlling miR expression remain poorly understood.

This study aims to explore the upstream regulators of miR-29a expression, shedding light on how these small RNA molecules are governed at the transcriptional level. miR-29a is a member of the miR-29 family of microRNAs, which play a crucial role in regulating various biological processes, including cell differentiation, proliferation, and apoptosis^2^. Highly expressed in the brain, miR-29 is linked to aging, metabolism, neuronal survival, and neurological disorders^3; 4; 5^. One study found that miR-29a is upregulated in the cortex and hippocampus during postnatal cerebrum development, where it is associated with neural activity through glutamate receptor activation, directly targets Doublecortin (DCX) to enhance axon branching, and is vital for neuronal development in mice^6^. Importantly, downregulation of miR-29a/b1 has been observed in neurodegenerative diseases such as Alzheimer’s^7^ and Huntington’s diseases^8^. Additionally, previous research showed that miR-29 levels in the blood serum of Parkinson’s disease patients were significantly reduced^9^. Mice lacking miR- 29a displayed signs of premature aging, including weight loss, decreased fat, muscle weakness, gait disturbances, and increased wrinkling^10^. Elevated levels of miR-29a in the human brain are associated with more rapid cognitive decline before death^11^. In rats, miR-29a knockout after Middle cerebral artery occlusion (MCAO) led to increased astrocyte proliferation and heightened glutamate release, exacerbating neurological damage^12^. Modulating miR-29a may contribute to neuroinflammatory responses and neuronal cell death, making it a potential therapeutic target for reducing neurodegeneration and preserving cognitive function.

To investigate the transcriptional regulation of miR-29a during neurogenesis, we utilized the Collaborative Cross (CC) mouse model, a resource designed to enhance genetic diversity and reproducibility in trait analysis for biomedical research^13; 14^. CC mice are derived from the genetic recombination of eight founder strains (five classical and three wild-derived), creating a genetically diverse population that mirrors the complexity of natural populations. This model captures over 90% of the genetic diversity in mouse species, providing exceptional resolution for genetic mapping and facilitating the identification of genes governing complex traits, such as disease susceptibility and neurodevelopmental processes.

The CC initiative is instrumental in studying intricate molecular mechanisms, enabling detailed exploration of transcriptional networks that regulate miRs. For instance, previous work in our lab demonstrated the power of CC mice in identifying upstream regulators of miR-9 during neurogenesis, uncovering key modulators that shape miR expression patterns critical for neuronal differentiation^15; 16^. Building on these findings, our current study focuses on miR-29a, leveraging the genetic diversity and mapping precision of CC mice to dissect the upstream regulatory elements influencing its expression. This approach aims to shed light on the molecular interactions and regulatory networks underlying miR-29a’s role in neuronal differentiation.

## Results

### Mapping regulatory genes from miR-29a expression profiling using CC mice strains

The goal of this experiment was to identify genes that regulate the expression of miR-29a in the hippocampus. Given the crucial role of miR-29a in neuronal development and its association with neurodegenerative diseases, understanding the mechanisms that control its expression is important. By utilizing expression profiling across genetically diverse Collaborative Cross (CC) mouse strains and performing Quantitative Trait Loci (QTL) analysis, the study aimed to locate genetic loci that influence miR-29a levels. The expression of miR-29a was examined in the hippocampi of male mice from 54 CC strains from the University of Tel Aviv, Israel, and Geniad, Australia (**Table S1**). The analysis revealed differential miR-29a expression among the CC strains, indicating genetic factors may affect miR-29a levels (**Figure 1A**). Next, the miR-29a expression data was processed with Gene Miner software to identify QTLs at specific genetic locations. A significant QTL for miR-29a expression was detected on chromosome 7, with a logarithm of the odds (LOD) score exceeding 12.5, suggesting a strong genetic association (**Figure 1B**). Ten candidate genes were identified within this QTL. Gene Ontology (GO) analysis of these genes showed enrichment in processes related to chromatin structure regulation and neuronal activity (**Figure 1C**). qPCR confirmed successful knockdown of candidate genes by siRNA, achieving over 70% reduction (**Figure 1D**, Oligonucleotides used in this study is mentioned in **Table S2**). Upon RNAi-mediated knockdown, Psmd13 and Nap1l4 demonstrated a significant increase in miR-29a expression compared to scrambled control (**Figure 1E**). These findings suggest that Psmd13 and Nap1l4, identified through QTL mapping on chromosome 7, are likely upstream regulators of miR-29a.

**Figure 1.**
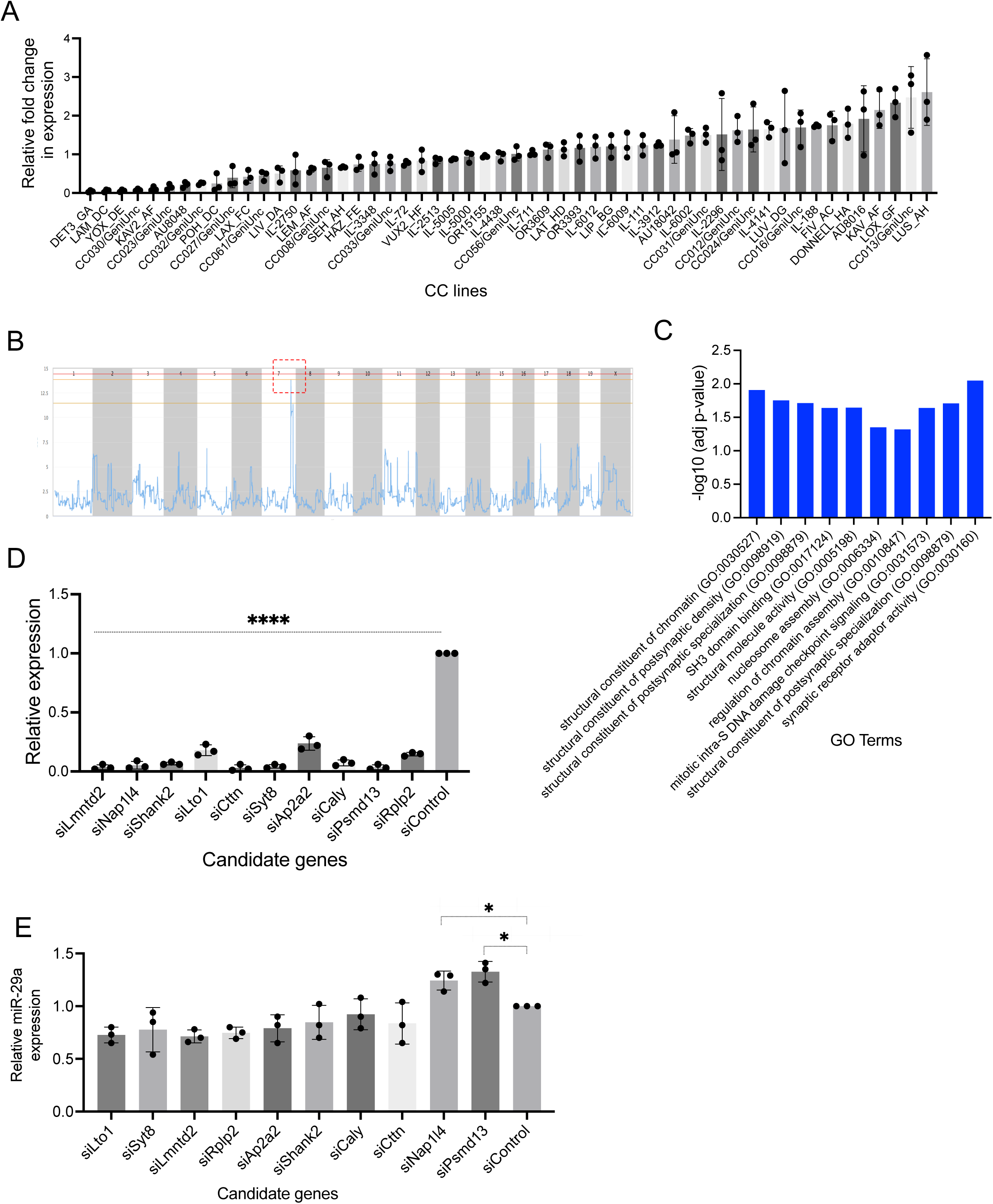
miR expression profiling and QTL analysis reveal upstream candidate genes of miR-29a. (A) Expression of miR-29a in hippocampi of 54 CC mice strains. (B) LOD score plot of miR-29a expression using Gene Miner. The QTL was detected on chromosome, Chr7 (red) with a significant score above 12.5. (C) Gene ontology (GO) analysis significantly represented for the candidate genes that were enriched in modules related to both regulation of chromatin structure and neuronal activities. Bar graphs indicate the statistical significance of the enrichment, as -log10 (adj p-value; cut-off level for significance, *p<0.05, adjusted by Benjamini-Hochberg correction). (D) The effectiveness of candidate gene knockdown using siRNAs was assessed through qPCR, comparing expression levels to those of the scrambled negative control. N= 3 experiments, mean ± SD, ****p<0.0001, One-way Anova. (E) Bar graph illustrating change in miR-29a expression, determined through RNAi and qPCR. Data from three individual experiments are shown here represented as mean ± SD.

### Screening of upstream candidate genes using neuronal differentiation assay in mNPCs

The objective was to determine if the knockdown of Psmd13 and Nap1l4 affects the differentiation of mNPCs and whether miR-29a is a key mediator in this process. Since miR-29a is known to play a role in neuronal development, the experiment aimed to assess whether Psmd13 and Nap1l4 knockdown influences the proportion of βIII-tubulin-positive neurons, a marker of neuronal differentiation, and whether this effect involves miR-29a. To perform this, mNPCs were isolated from mouse hippocampus and treated with siRNAs targeting Psmd13 and Nap1l4 (siPsmd13 and siNap1l4), followed by a three-day differentiation period. Quantification using the βIII- tubulin marker revealed that Psmd13 knockdown significantly increased the percentage of βIII-tubulin-positive cells compared to the scrambled siRNA control (**Figures 2 A-B**), and this was corroborated by flow cytometry analysis (**Figures 2C-D**). To validate these findings, rescue experiments were performed by overexpressing Psmd13, which reduced neuronal differentiation (**Figure S1 A-B**). Additionally, inhibition of miR-29a in mNPCs resulted in a notable increase in neuronal differentiation, as shown by the higher number of βIII-tubulin-positive cells in both immunostaining (**Figures 2 E-F**) and flow cytometry (**Figures 2 G-H**). Overexpression of miR-29a using miR-29a mimics decreased neuronal differentiation (**Figure S1 C-D**). When Psmd13 knockdown was combined with miR- 29a mimics, there was a significant reduction in the number of βIII-tubulin-positive cells compared to Psmd13 knockdown alone (**Figures 2 I-J**), confirmed by flow cytometry (**Figures 2 K-L**). These findings indicate that Psmd13 regulates neuronal differentiation, and miR-29a likely mediates Psmd13’s effects on this process.

**Figure 2.**
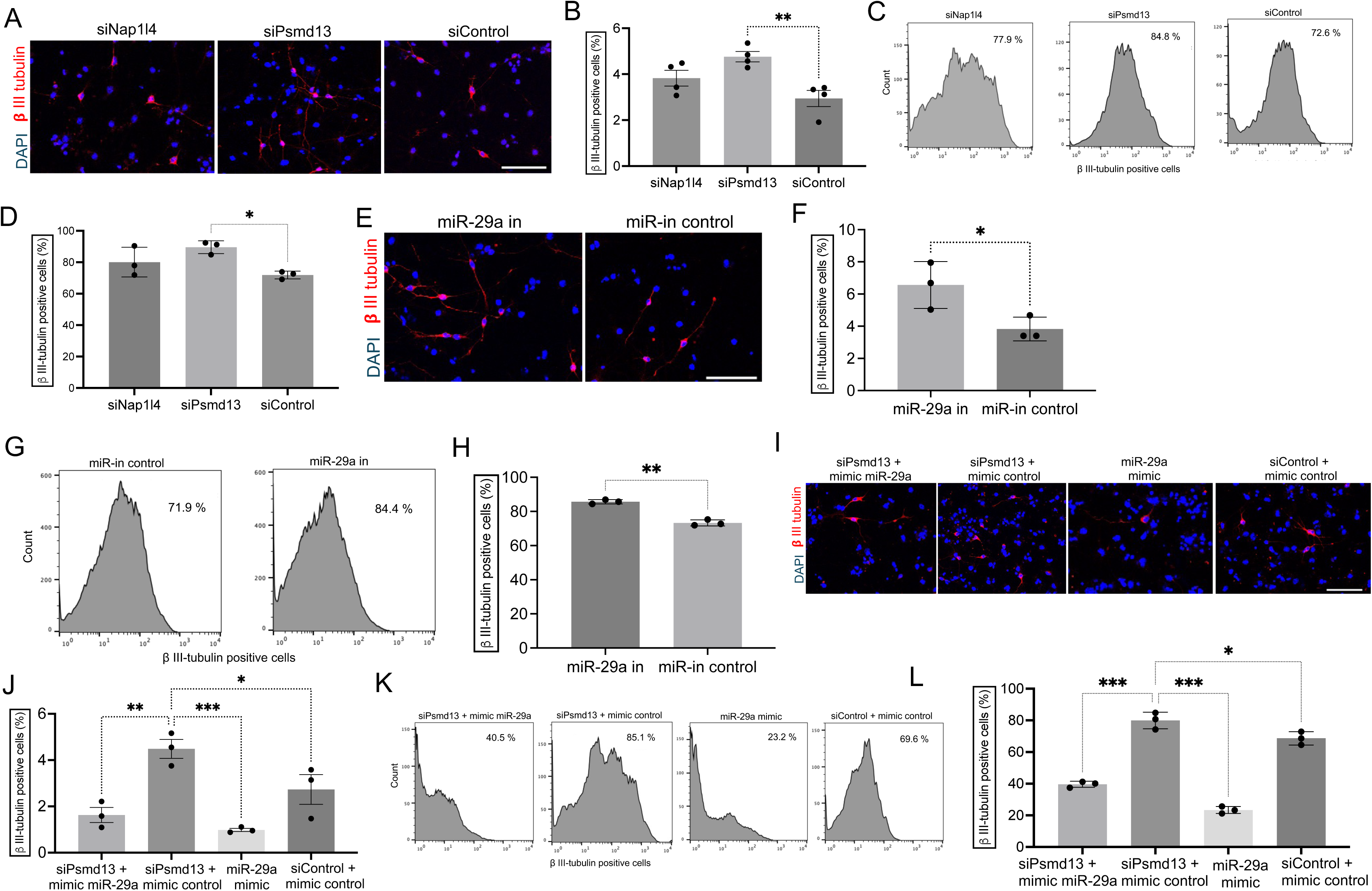
Screening of upstream candidate genes using neuronal differentiation in mNPCs. (A-B) Role of candidate genes in neuronal differentiation. Representative images (A) and quantification (B) of βIII-tubulin positive cells using immunostaining in differentiated mNPCs transfected with individual siRNAs and scrambled siRNA negative control. N= 4 experiments. mean ± SD, **p<0.01. One-way Anova. Image scale bars=10 μm. (C-D) Representative histograms (C) and quantification (D) of βIII-tubulin detection using flow cytometry in differentiated mNPCs transfected with siPsmd13, siNap1l4 and siRNA of non-targeting control. N= 3 experiments. mean ± SD, *p<0.05. One-way Anova. (E-F) miR-29a alter neuronal differentiation of mNPCs. Representative images (E) and quantification (F) of βIII-tubulin positive cells using immunostaining in differentiated mNPCs transfected with miR-29a inhibitor and miR negative control. N=3 experiments. mean ± SD, *p<0.05. Unpaired T-test. Image scale bars=10 μm. (G-H) Representative histograms (G) and quantification (H) of βIII-tubulin detection using flow cytometry in differentiated mNPCs transfected with miR-29a inhibitor and inhibitor control. N= 3 experiments. mean ± SD, **p<0.01. Unpaired T-test. (I-J) Psmd13 acts through miR-29a to alter neuronal differentiation. Representative images (I) and quantification (J) of βIII-tubulin positive cells using immunostaining in differentiated mNPCs transfected with siRNA non-targeting control, siPsmd13 plus mimic control, siPsmd13 plus miR-29a mimics and miR-29a mimics. N= 3 experiments. mean ± SD, **p<0.01, *p<0.05. One-way Anova. Image scale bars=10 μm. (K-L) Representative histograms (K) and quantification (L) of βIII-tubulin detection using flow cytometry in differentiated mNPCs transfected with control plus siRNA non-targeting control, siPsmd13 plus mimic control siPsmd13 plus miR-29a mimics and miR-29a mimics. N= 3 experiments. mean ± SD, *p<0.05. One-way Anova.

### Psmd13 associates with Dicer to regulate miR-29a expression in mNPCs

The aim of this experiment was to explore the relationship between Psmd13 and Dicer in mNPCs and how this interaction influences the expression of miR-29a, particularly in the context of neuronal differentiation. By examining the effects of Psmd13 knockdown on miR-29a expression through qPCR, a significant increase in miR-29a levels was observed in Psmd13-depleted mNPCs in the undifferentiated state (**Figure 3A**), but a notable reduction was seen after differentiation (**Figure 3B**). This suggests that Psmd13 regulates miR-29a, maintaining its expression throughout neuronal differentiation. Additionally, Dicer expression, was analysed and showed a significant rise in miR-29a levels in undifferentiated Psmd13-depleted mNPCs (**Figure 3C**) but decreased after differentiation (**Figure 3D**), indicating that Psmd13 regulates Dicer. These changes were corroborated by western blot analysis, which demonstrated similar trends in Dicer protein levels (**Figures 3 E-F and Figures S2 A-B**). To evaluate the functional consequences of Psmd13 regulation of miR-29a, dual luciferase reporter assays were performed. We utilized both the miR- 29a promoter sequence and a mutant version of the miR-9 promoter sequence in the 3’ UTR region of the luciferase reporter gene. After transfecting mNPCs with siRNAs targeting Psmd13, along with vectors containing miR-29a and mutant miR-29a reporter genes, we assessed luciferase activity. These assays showed that miR-29a effectively regulates Psmd13 expression, with increased activity in undifferentiated cells (**Figure 3G**) and decreased activity in differentiated cells (**Figure 3H**). However, the luciferase activity of the vector harboring the mutant miR-29a remained unchanged. These findings confirm the functional role of miR-29a in neuronal differentiation and highlight Psmd13’s involvement in regulating miR-29a. Both Psmd13 and Dicer were localised in the cytoplasm in the undifferentiated mNPCs but showed cytoplasmic and nuclear localisation in the differentiated state (**Figures S2 C-D**). Further investigation of the interaction between Psmd13 and Dicer through co-immunoprecipitation (Co-IP) and reverse Co-IP experiments revealed that Psmd13 physically associates with Dicer in undifferentiated mNPCs (**Figures 3 I-J and Figures S2 E-F**). However, this interaction was significantly diminished in differentiated cells (**Figures 3 K-L and Figures S2 G-H**), suggesting that the Psmd13-Dicer interaction is differentiation-dependent. This differentiation-dependent dynamic may play a critical role in regulating miR-29a regulation.

**Figure 3.**
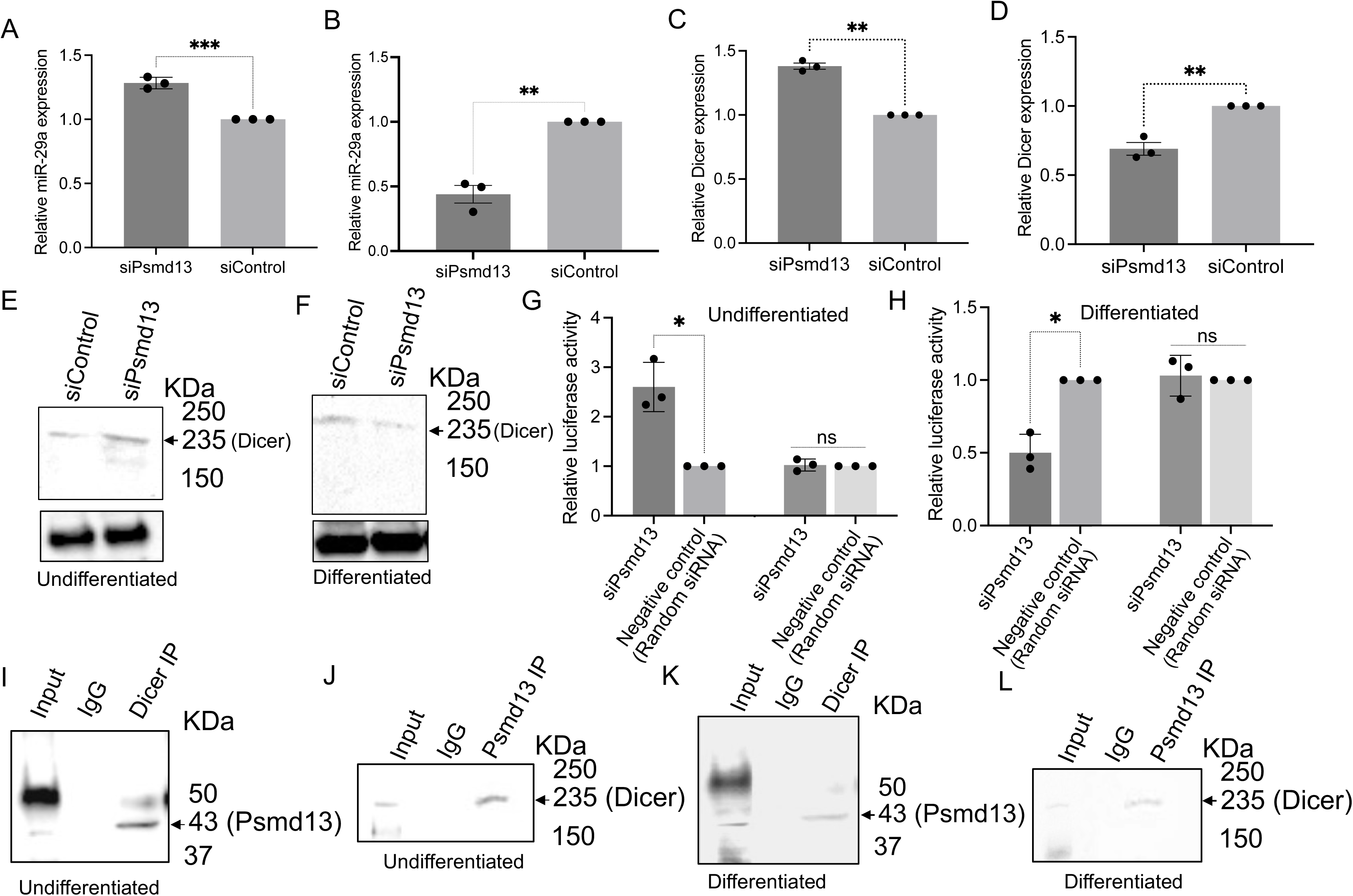
Psmd13 associates with Dicer and regulate miR-29a expression in mNPCs. (A) Quantitative analysis of the expression fold change of miR-29a (by Taqman PCR), in siControl and siPsmd13 undifferentiated mNPCs. (B) Quantitative analysis of the expression fold change of miR-29a (by Taqman PCR), in siControl and siPsmd13 differentiated mNPCs. (C) Quantitative analysis of the expression fold change of Dicer in siControl and siPsmd13 mNPCs in undifferentiated mNPCs. (D) Quantitative analysis of the expression fold change of Dicer in siControl and siPsmd13 mNPCs in differentiated mNPCs. N= 3 experiments. mean ± SD, **p<0.01. Unpaired T-test. (E) Western blotting was conducted using specific antibodies to detect Dicer and Actin proteins in extracts from control and Psmd13-depleted mouse neural progenitor cells under undifferentiated conditions. (F) Western blotting was conducted using specific antibodies to detect Dicer and Actin proteins in extracts from control and Psmd13-depleted mouse neural progenitor cells under differentiated conditions. (G) Dual luciferase reporter assay in mNPCs for miR-29a and mutated miR-29a sites. Values for Firefly Luciferase luminescence normalized to Renilla luminescence. Fold change relative to control in undifferentiated mNPCs. (H) Dual luciferase reporter assay in mNPCs for miR-29a. Values for Firefly Luciferase luminescence normalized to Renilla luminescence. Fold change relative to control in differentiated mNPCs. N= 3 experiments. mean ± SD, *p<0.05., ns – not significant. Unpaired T-test. (I) Co-IP of Dicer was performed with corresponding antibody followed by western blotting to detect endogenous Psmd13 proteins in undifferentiated mNPCs. Input and IgG antibody was used as controls for the experiment. (J) Co-IP of Psmd13 was performed with corresponding antibody followed by western blotting to detect endogenous Dicer proteins in undifferentiated mNPCs. Input and IgG antibody was used as controls for the experiment. (K) Co-IP of Dicer was performed with corresponding antibody followed by western blotting to detect endogenous Psmd13 proteins in differentiated mNPCs. Input and IgG antibody was used as controls for the experiment. (L) Co-IP of Psmd13 was performed with corresponding antibody followed by western blotting to detect endogenous Dicer proteins in differentiated mNPCs. Input and IgG antibody was used as controls for the experiment.

### Psmd13 depletion impacts global miR regulation in mNPCs

We performed small RNA sequencing (sRNA-seq) to compare global miR expression profiles in control and Psmd13-depleted mNPCs in both undifferentiated and differentiated states. Two independent libraries were analyzed for siCtrl and siPsmd13 samples. Sequencing reads were mapped to unique sites in the mouse genome (mm10) using Bowtie (v2.3.5), and differential expression analysis was conducted with DESeq2. The analysis confirmed that miR-29a was upregulated in undifferentiated mNPCs but downregulated in differentiated mNPCs following Psmd13 depletion compared to controls (**Figures 4A-B**).

**Figure 4.**
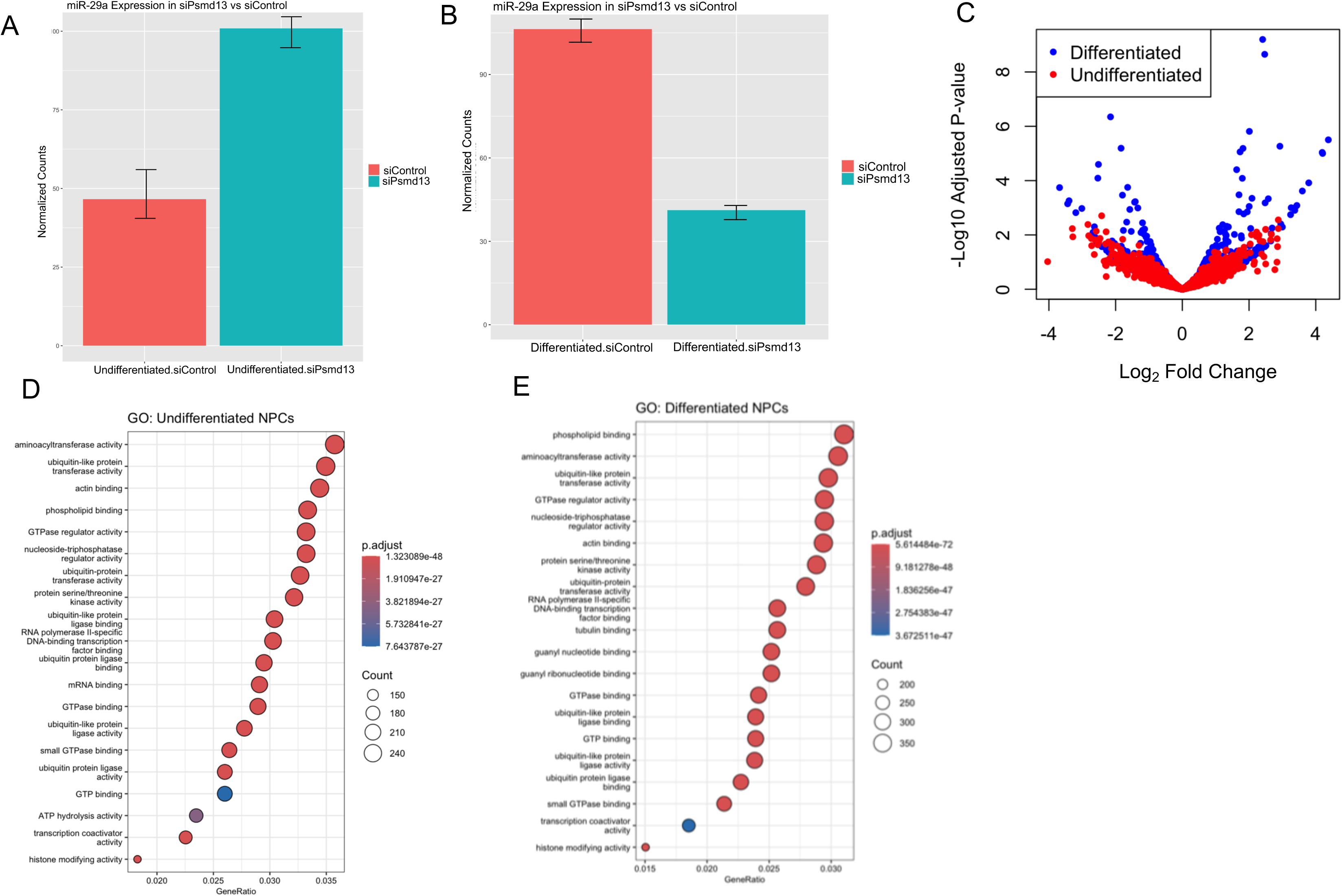
Psmd13 depletion impacts global miR regulation in mNPCs. (A) Box plots of read distributions for significantly differentially bound sites in the undifferentiated mNPCs between Psmd13-depleted and control groups. (B) Box plots of read distributions for significantly differentially bound sites in the differentiated mNPCs between Psmd13-depleted and control groups. (C) Volcano plot of differentially expressed genes (padj < 0.05). Log2 fold change between differentiated and undifferentiated mNPCs is plotted on the x-axis and the −log10 of the padj value is plotted on the y-axis. (D) The top 20 enriched GO terms of molecular function in the undifferentiated state. Circle size indicates the number of genes enriched in each term. Color saturation represents the significance level. (E) The top 20 enriched GO terms of molecular function in the differentiated state. Circle size indicates the number of genes enriched in each term. Color saturation represents the significance level.

A volcano plot revealed more differentially expressed miRs in the differentiated state compared to the undifferentiated state after Psmd13 knockdown (**Figure 4C**). **Table S7** showed that over 1,000 miRs were differentially expressed in undifferentiated and differentiated mNPCs after Psmd13 depletion. To assess the functional impact of these changes, we performed GO enrichment analysis using enrichGO. GO analysis of differentially expressed miRs (DEMs) revealed significant enrichment of terms associated with ubiquitin processes, transcription coactivator activity, RNA polymerase-specific DNA binding, and histone modification. Additionally, tubulin-binding genes were enriched in differentiated mNPCs (**Figures 4D-E**).

To further explore miR-gene interactions after Psmd13 depletion, we constructed interaction networks using Multimir. Interestingly, siPsmd13 showed an increased number of interactions in the differentiated state compared to the undifferentiated state (**Figures S3A-B**). These findings confirm that Psmd13 depletion significantly impacts global miR regulation in both differentiated and undifferentiated mNPCs.

### Psmd13 dependency for Dicer binding in mNPCs

The objective of this study was to investigate the co-binding patterns of Psmd13 and Dicer and assess their functional co-regulation during neuronal differentiation. We hypothesized that Psmd13, a proteasome subunit, physically interacts with Dicer and that this interaction influences key regulatory pathways, such as miR-29a regulation, during neuronal differentiation.

Using chromatin immunoprecipitation sequencing (ChIP-seq), we aimed to explore whether Psmd13 and Dicer co-localize at specific genomic loci, including those involved in neuronal differentiation, and how their binding dynamics change between undifferentiated and differentiated mNPCs. ChIP-seq density heatmaps and global binding profiles revealed substantial co-binding of Psmd13 and Dicer at numerous loci in mNPCs, with enrichment around 5,000-bp regions flanking transcription start sites (TSS). This suggests that Psmd13 and Dicer may play a role in transcriptional regulation at these sites. Integrating histone ChIP-seq data (obtained from the publicly available datasets - ENCFF746JVZ, ENCFF221ABQ, ENCFF095VNG, ENCFF787DWM, ENCFF740AHK, ENCFF902SQA and ENCFF955DGO) with our ChIP-seq profiles further indicated that these co-bound sites associate with histone modifications indicating transcriptional regulation (**Figures 5 A-B**).

**Figure 5.**
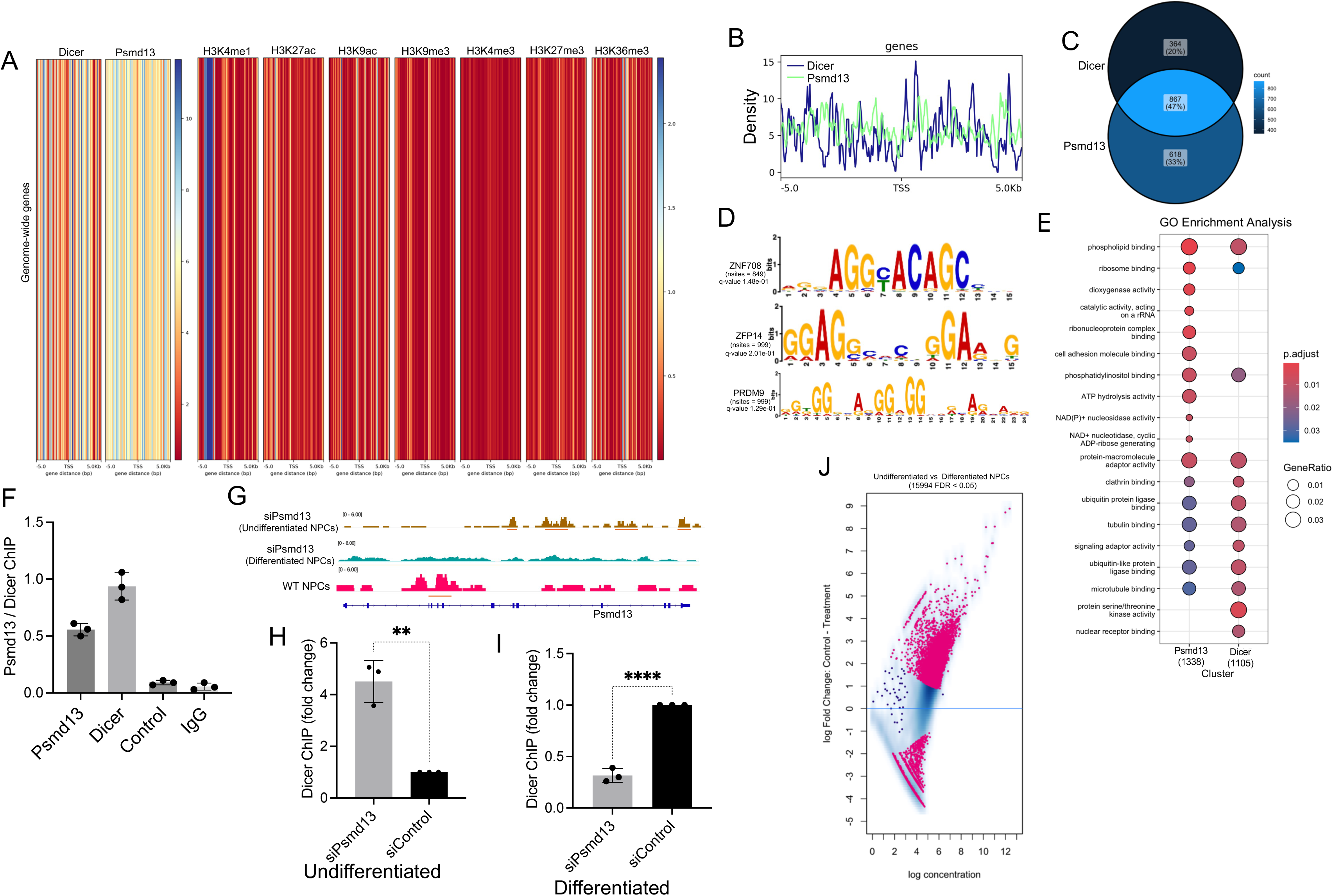
Psmd13 dependency for Dicer binding at miR-29a locus in mNPCs. (A) ChIP-seq density heatmaps of Psmd13 and Dicer binding, and ChIP seq of different histone marks in mNPCs are shown within the extended gene regions (−5 kb of TSS to +5 kb of TSS) of total genome-wide genes. (B) Density map showing the global binding profiles of Dicer and Psmd13 ChIP signal. The x axis represents the distance to the TSS. The y axis represents per base read coverage. (C) Overlap of the genes annotated between Psmd13 and Dicer displayed as Venn diagram. The P-value was calculated using Fisher’s exact test. (D) Transcription factor binding motifs enriched in the Dicer and Psmd13 ChIP-seq peak dataset identified by MEME-ChIP. (E) The top 20 enriched GO terms of molecular function. Circle size indicates the number of genes enriched in each term. Color saturation represents the significance level. (F) Individual ChIP analysis showing chromatin co-localization of Psmd13 and Dicer at the miR-29a genomic loci in mNPCs (N = 3, data are shown as mean ± SD). The binding was confirmed against a control region on the miR-29a locus. (G) ChIP-seq profiles showing Dicer binding at Psmd13 loci in Psmd13-depleted undifferentiated and differentiated mNPCs, as well as in WT mNPCs. Peaks are visualized in IGV, with undifferentiated samples shown in brown, differentiated in green, and WT in magenta, all exhibiting enrichment over the Input control. WT = wild-type (H) Individual ChIP analysis comparing chromatin binding of Dicer at miR-29a genomic loci in control and Psmd13-depleted undifferentiated mNPCs (N = 3, data are shown as mean ± SD). Unpaired t test was used for statistical analysis (**p < 0.01). (I) Individual ChIP analysis comparing chromatin binding of Dicer at miR-29a genomic loci in control and Psmd13-depleted differentiated mNPCs (N = 3, data are shown as mean ± SD). Unpaired t test was used for statistical analysis (****p < 0.0001). (J) Fold changes of Dicer binding versus signal intensity between undifferentiated and differentiated mNPCs are visualized as MA plot. Pink represents differentially bound peaks (FDR < 0.05). The x axis values (“log concentration”) represent logarithmically transformed, normalized counts, averaged for all samples, for each site. The y axis values represent log 2 (fold change) values.

A Venn diagram displayed in **Figure 5C** shows that 47% of genes bound by Psmd13 and Dicer exhibit statistically significant co-binding. The bar plot in **Figure S4 A** illustrates the distribution of Psmd13, Dicer, and histone mark peaks relative to TSS, while **Figure S4 B** shows the percentage of annotated features associated with each peak type. Additionally, the metaplot in **Figure S4 C** shows the normalized binding intensity of Psmd13, Dicer, and histone marks across ±5 kb regions around TSS. Motif analysis using MEME-ChIP identified three transcription factor binding motifs— ZNF708, ZFP14, and Prdm9—enriched in the Dicer and Psmd13 ChIP-seq peak dataset. These transcription factors are known to regulate neuronal differentiation and chromatin remodeling ^17; 18; 19^, suggesting that Psmd13 and Dicer may cooperate with these factors to modulate gene expression during neuronal differentiation (**Figure 5D**). Gene ontology (GO) analysis revealed that co-bound regions were enriched for terms related to protein binding and enzymatic activity, implicating roles in transcription regulation, RNA processing, and signalling pathways, and indicating a complex interplay between Dicer activity, miR expression, and proteasomal regulation in neuronal differentiation (**Figure 5E**).

Regarding Dicer’s interaction with Psmd13, ChIP-seq and individual ChIP analyses showed baseline Dicer binding at Psmd13 loci, potentially to regulate miR-29a in WT NPCs. In Psmd13-depleted mNPCs, Dicer binding at Psmd13 increased in the undifferentiated state but was absent in the differentiated state. This loss of Dicer binding during differentiation suggests that Psmd13 may be required to maintain Dicer occupancy at the loci in differentiated mNPCs (**Figures 5 F-G**). Additionally, Psmd13 depletion led to increased Dicer binding at miR-29a loci in undifferentiated mNPCs but decreased binding in differentiated cells as determined by ChIP-PCR (**Figures 5 H-I**).

A detailed comparison of Dicer binding between undifferentiated and differentiated mNPCs, visualized via IGV tracks and MA plots, confirmed that Dicer binding at Psmd13 loci is significantly higher in the undifferentiated state. Fold-change analysis (MA plots) indicated that several Dicer-bound peaks are differentially enriched during differentiation (FDR < 0.05), with significantly differentially bound regions marked in pink. Box plot analysis revealed differences in binding patterns, as indicated by variations in read distributions between differentiated and undifferentiated mNPCs. (**Figure 5J** and **Figures S4 D-E**). In order to determine the co-localisation of Psmd13 and Dicer at miR loci, we identified miRs critical for neuronal related processes and determined if Psmd13 and Dicer co-localise in the specific miR loci. Moreover, we compared the co-binding to the fold change expression as determined by the sRNA-seq. A total of 37 miRs were tested in this category including the differentiated and undifferentiated ChIP-seq datasets. We found that Psmd13 bound to 13 (81%) of 16 Dicer-bound miR loci, indicative of a significant overlap of Dicer/Psmd13 binding at miR loci (**Table S8**).

Together, these data demonstrate that Psmd13 and Dicer bind to miR-29a genomic locus, modulating miR-29a expression and influencing differentiation states in mNPCs.

### Impact of proteasome inhibition on Dicer dynamics and miR-29a regulation in mNPCs

The 26S proteasome is a complex assembly composed of a 20S core particle and one or two 19S regulatory particles, responsible for selectively degrading ubiquitin-tagged proteins within cells. This intricate structure plays a vital role in cellular homeostasis by facilitating the recognition, unfolding, and translocation of substrates into the catalytic core, thereby regulating protein turnover and modulating various signalling pathways^20^. Psmd13, a non-ATPase regulatory subunit of the 26S proteasome, plays a crucial role in the recognition and processing of substrates, facilitating their degradation and influencing cellular responses to stress and signalling pathways^21^ (**Figure S5 A**). To assess the effects of proteasome inhibition on Dicer levels in mNPCs, we initially evaluated the expression of the proteasomal subunit Psmd13. Western blot analysis showed that treatment with 5 μM MG132 (proteasome inhibitor) resulted in a time-dependent decrease in Psmd13 protein levels compared to the DMSO control, whereas 0.2 μM Capzimin (proteasome inhibitor) did not affect its expression (**Figure 6A** and **Figure S5 B**). Quantitative analysis of Psmd13 mRNA revealed a significant reduction in expression following MG132 treatment, particularly at 12 and 20 hours (**Figure 6B**). Next, we examined Dicer protein levels, which were significantly decreased in MG132-treated mNPCs upon long exposure (**Figure 6C** and **Figure S5 C**). This finding was supported by quantitative analysis of Dicer mRNA, which also showed a significant time-dependent decline in response to MG132 treatment (**Figure 6D**). Additionally, the expression of miR-29a, a miR potentially regulated by Dicer, was measured using Taqman PCR, revealing a notable downregulation over time in MG132-treated mNPCs (**Figure 6E**). Assessment of proteasomal activity following MG132 treatment indicated a marked decrease with prolonged exposure (**Figure S5 D**). To explore the functional consequences of proteasome inhibition, we evaluated mNPC differentiation into βIII-tubulin positive neurons. Immunostaining demonstrated a significant reduction in the number of βIII-tubulin positive cells in differentiated MG132-treated mNPCs compared to control and Psmd13-depleted mNPCs (**Figures 6 F-G**). Flow cytometry analysis confirmed these findings, showing a decrease in βIII-tubulin positive cells among MG132-treated, untreated, and non-targeting control mNPCs (**Figures S5 E-F**). To elucidate the underlying molecular mechanisms, we conducted ChIP-seq analysis to examine Dicer binding at Psmd13 loci in both MG132-treated and untreated mNPCs. The ChIP-seq profiles indicated distinct binding patterns, with decreased enrichment observed at specific loci in MG132- treated cells (**Figure 6H**). Further analysis revealed that Dicer interacted with miR- 29a genomic loci across MG132, Psmd13-depleted, and control mNPCs (**Figure 6I**). An MA plot visualized the fold changes in Dicer binding between MG132-treated and untreated mNPCs, highlighting differentially bound peaks (FDR < 0.05) (**Figure 6J**). This analysis illustrated a shift in binding dynamics, underscoring the regulatory role of Dicer in response to proteasome inhibition. Box plots further illustrated differences in Dicer binding at significantly differentially bound sites (**Figure 6K**), while heatmaps of Dicer read enrichments revealed pronounced differences between MG132-treated and untreated mNPCs within a 1500 bp window centered around transcription start sites (TSS) (**Figure 6L**). Gene ontology (GO) analysis of co-bound regions demonstrated significant enrichment for terms related to chromatin structure regulation and neuronal activities, The top 20 GO terms included molecular functions such as chromatin binding, transcription regulation, and synapse organization, emphasizing the relevance of these interactions to neurodevelopment (**Figure S5 G**). Additionally, a Venn diagram illustrating the overlap of genes between the MG132-treated and untreated ChIP-seq profiles highlighted key differences in gene regulatory networks (**Figure S5 H**). Overall, these findings suggest that proteasome inhibition leads to the accumulation of regulatory proteins, including Dicer, which is essential for miR regulation; this, in turn, results in the time-dependent downregulation of miR-29a in mNPCs, indicating that proteasomal activity is crucial for maintaining miR balance and significantly influencing neuronal differentiation.

**Figure 6.**
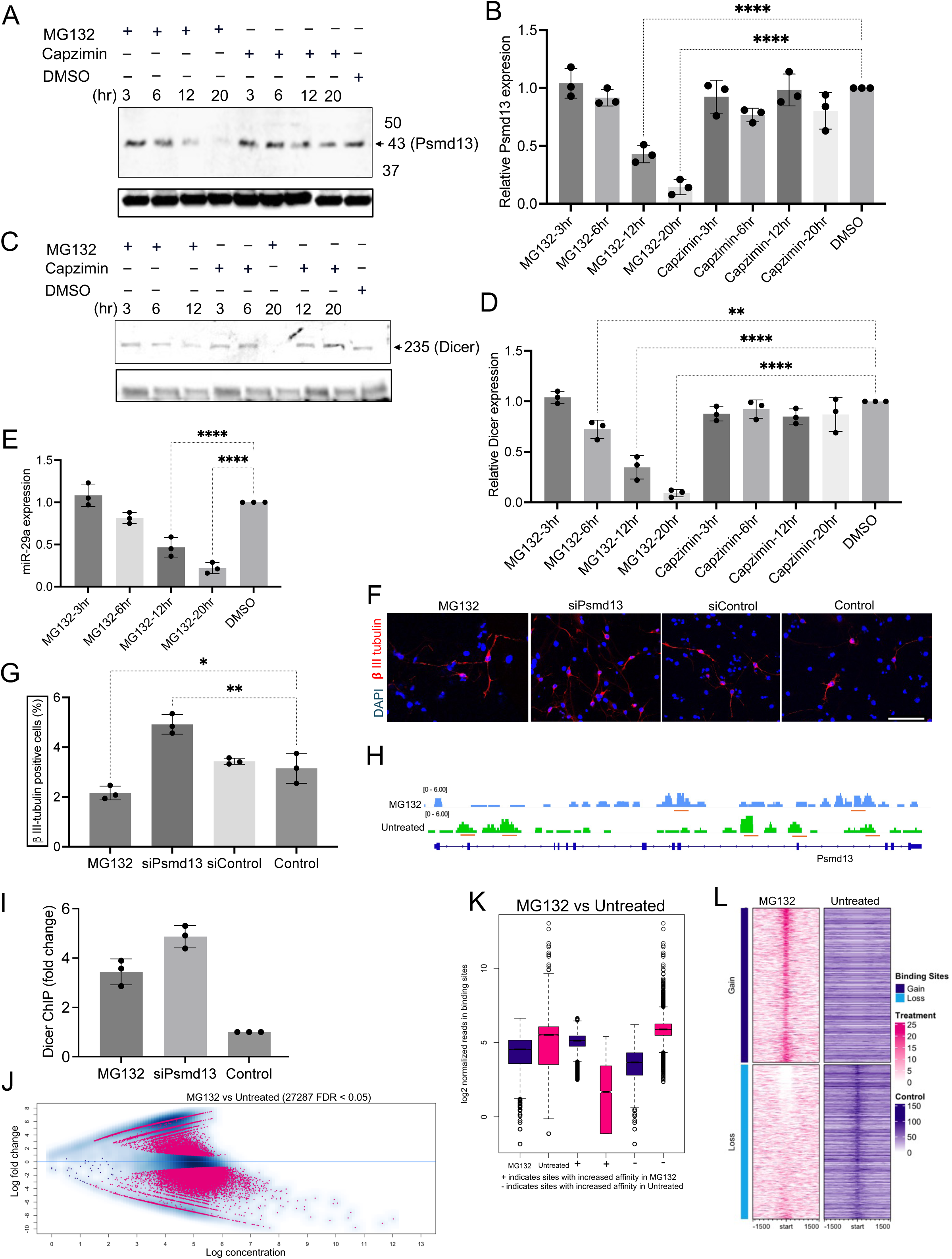
Effect of proteasome inhibition on Dicer levels and miR-29a Expression in mNPCs. (A) Western blot showing PSMD13 protein levels in mNPCs following treatment with 5 μM MG132, 0.2 μM Capzimin, and DMSO control for the specified durations. B-actin was used as a loading control. (B) Quantitative analysis of the expression fold change of Psmd13 mRNA in MG132 treated mNPCs in a time-dependant experiment (normalised to control). N= 3 experiments. mean ± SD, ****p<0.0001. One-way Anova. (C) Western blot showing the levels of Dicer protein after treatment with 5 μM MG132, 0.2 μM Capzimin and DMSO control in mNPCs for the indicated time duration. B-actin was used as a loading control. (D) Quantitative analysis of the expression fold change of Psmd13 mRNA in MG132 treated mNPCs in a time-dependant experiment (normalised to control). N= 3 experiments. mean ± SD, **p<0.01, ****p<0.0001. One-way Anova. (E) Quantitative analysis of the expression fold change of miR-29a (by Taqman PCR), in MG132 treated mNPCs in a time-dependant experiment (normalised to control). N= 3 experiments. mean ± SD, ****p<0.0001. One-way Anova. (F-G) Representative images (F) and quantification (G) of βIII-tubulin positive cells using immunostaining in differentiated mNPCs treated with MG132, Psmd13-depleted and control mNPCs. N= 3 experiments. mean ± SD, **p<0.01, *p<0.05. One-way Anova. Image scale bars=10 μm. (H) ChIP-seq profiles showing Dicer binding at Psmd13 loci in MG132 treated and untreated mNPCs. Peaks are visualized in IGV, with MG132 shown in blue and untreated in green, all exhibiting enrichment over the Input control. (I) Individual ChIP analysis comparing chromatin binding of Dicer at miR-29a genomic loci in MG132, Psmd13-depleted and control mNPCs (N = 3, data are shown as mean ± SD). (J) Fold changes of Dicer binding between MG132 and untreated mNPCs are visualized as MA plot. Pink represents differentially bound peaks (FDR < 0.05). The x axis values (“log concentration”) represent logarithmically transformed, normalized counts, averaged for all samples, for each site. The y axis values represent log 2 (fold change) values. (K) Box plots of read distributions for significantly differentially bound sites in the MG132 and untreated mNPCs. (L) Heatmaps showing the enrichment of Dicer reads between MG132 and untreated mNPCs in a 1500 bp window centered around the TSS. Scale is as indicated in the signal.

## Discussion

Based on our findings, we propose a working model for miR-29a expression regulation in mNPCs (**Figure 7**). This study provides significant insights into the genetic and molecular regulation of miR-29a expression and its role in neuronal differentiation, leveraging the genetic diversity of Collaborative Cross (CC) mouse strains to map regulatory genes. miR-29a, a miR implicated in neuronal development and neurodegeneration, was observed to vary in expression across different CC strains, pointing to genetic factors that modulate its levels in the hippocampus. Through quantitative trait loci (QTL) analysis, a critical locus on chromosome 7 was identified with a high LOD score, suggesting a strong genetic association. Ten candidate genes within this locus, particularly Psmd13 and Nap1l4, were identified as potential regulators of miR-29a expression.

**Figure 7.**
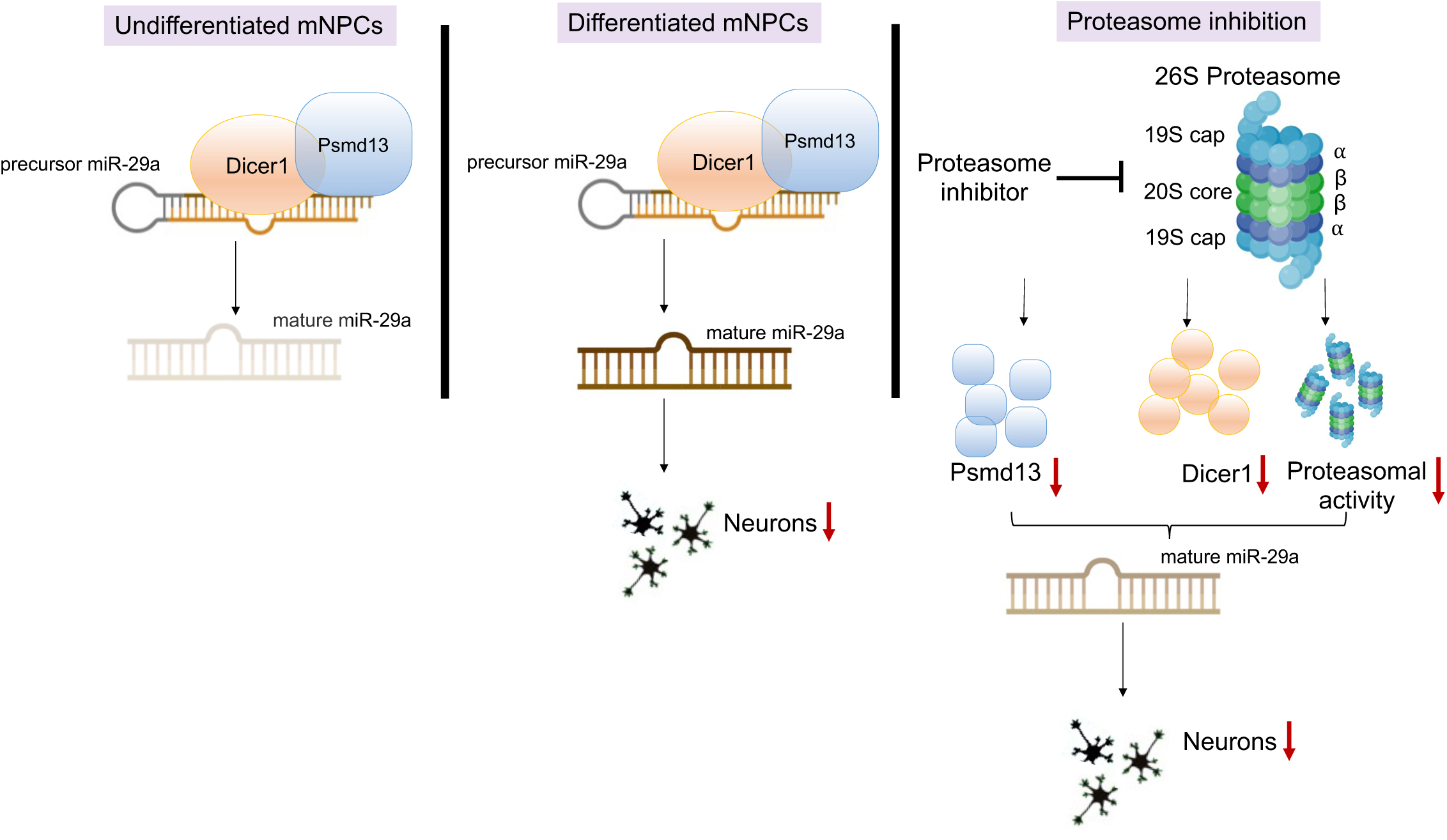
Psmd13-Dicer interaction modulates miR-29a expression and neuronal differentiation in mNPCs. In the undifferentiated state, Psmd13 co-localizes with Dicer in the cytoplasm and helps regulate miR-29a expression. Upon Psmd13 knockdown, both Dicer and miR-29a levels increase. Elevated miR-29a levels further reinforce the repression of differentiation-associated genes, preserving the progenitor state. In the differentiated state, inhibition of Psmd13 and miR-29a enhances neuronal differentiation, suggesting that miR-29a acts downstream of Psmd13 to suppress differentiation by targeting genes that promote this process. Proteasome inhibition disrupts the Psmd13-Dicer interaction, reducing miR-29a levels and impairing differentiation. Changes in miR-29a levels are visually represented by shifts in its color (brown).

### Psmd13 and miR-29a: A Regulatory Axis in Neuronal Differentiation

The hallmarks of neurodegenerative disorders in general are neuronal inclusions of misfolded proteins, which often contain ubiquitylated proteins and proteasomes. The 26S proteasome is a multi-subunit protease complex critical for protein quality control, synaptic plasticity, and neuronal function. Neurodegenerative disorders are caused by the dysfunction of 26S Proteasomes. In mice with brain region-specific knockout of a 19S subunit, neurodegeneration is accompanied by accumulation of ubiquitylated α-synuclein inclusion ^22^. Psmd13 is a non-enzymatic component of the 26S proteasome that influences proteasomal function. PSMD13 gene silencing suppressed the production of proinflammatory mediators by modulating ubiquitin-proteasome system-mediated neuroinflammation ^21^.

During differentiation, Psmd13 helps maintain the expression of Dicer and miR-29a. When Psmd13 is knocked down, Dicer and miR-29a levels drop, leading to enhanced neuronal differentiation. This suggests that miR-29a normally suppresses neuronal differentiation, and its reduction (due to decreased Dicer) allows more neurons to form. In undifferentiated cells, Psmd13 suppresses Dicer and miR-29a expression. When Psmd13 is knocked down, Dicer and miR-29a levels rise, indicating that Psmd13 negatively regulates them at this stage. This could mean that in the undifferentiated state, higher levels of miR-29a help maintain the cells in an undifferentiated state. As a result, Psmd13 acts as a regulator whose effects on Dicer and miR-29a vary depending on the cell’s developmental stage. Furthermore, miR-29a functions downstream of Psmd13 to repress differentiation, likely by targeting differentiation-promoting genes. The absence of both Psmd13 and miR-29a removes this repression, allowing cells to proceed with differentiation. Our findings are consistent with previous studies showing Dicer’s essential role in neurogenesis through miR regulation ^23^, and miR-29a’s role in promoting neuronal differentiation by repressing cell cycle-related genes ^24^ .

#### Psmd13-Dicer Interaction and miR-29a Regulation

Microarray analysis confirmed the presence of miRs in the samples of purified proteasomes both in the nuclear and cytoplasmic fractions ^25^. Psmd13 might have a predominantly cytoplasmic localization under basal conditions but may influence nuclear processes indirectly by interacting with proteins or complexes that shuttle between the cytoplasm and nucleus under specific processes. Our study also confirmed that Psmd13 is localized in the cytoplasm with Dicer, and this interaction may act as a chaperone to support Dicer’s function during differentiation, facilitating its shuttling between the nucleus and cytoplasm. Psmd13 depletion significantly alters global miR regulation in mNPCs, with distinct effects observed in differentiated and undifferentiated states, including changes in miR-29a expression and enrichment of pathways linked to transcription regulation, histone modification, and tubulin binding. Furthermore, Psmd13 and Dicer co-localize at chromatin regions, including miR-29a’s regulatory loci. Psmd13’s association with histone marks and transcription factor motifs (e.g., ZNF708, ZFP14, Prdm9) suggests that Psmd13 influences chromatin accessibility, likely enhancing transcription of differentiation genes. These results suggest Psmd13 not only stabilizes Dicer but may also regulate chromatin structure, facilitating miR-mediated gene regulation essential for neurogenesis. Thus, Psmd13 serves as a modulator of neuronal differentiation, influencing the balance between progenitor cell proliferation and differentiation through a feed-forward mechanism.

#### Proteasomal Activity and miR regulation

Proteasomal activity also emerged as a critical factor in regulating Dicer and miR- 29a levels. Proteasome inhibition with MG132 reduced Dicer and miR-29a levels, as well as neuronal differentiation, underscoring the proteasome’s role in stabilizing key components of miR regulation. Consistent with previous studies, proteasomes likely interact with RNAi machinery, including Dicer and Ago proteins, to regulate miR turnover and function ^26; 27^. Hsp90 and mTOR signalling may also influence this process by stabilizing proteasomes or mediating ubiquitination of miR regulation proteins, such as Drosha and Dicer through E3 ubiquitin ligase MDM2. Furthermore, ZFP14, a transcription factor identified in our study, interacts with MDM2 through its zinc finger domains, promoting MDM2’s ubiquitination ^18; 19^ which might explain the interaction between proteasomes with miR, which, in turn, binds the miR-processing complex, Dicer. Moreover, the disruption of Dicer binding patterns upon proteasome inhibition further supports a feedback mechanism linking proteasomal activity to miR regulation during neuronal differentiation.

#### Implications and Future Directions

Overall, this study identifies Psmd13 as a critical upstream regulator of miR-29a and demonstrates its involvement in neuronal differentiation through interactions with Dicer. Thus, it will be interesting for future studies to investigate whether Psmd13 and proteasome pathway play a conserved role in co-transcriptional miR-29a regulation and to elucidate additional regulatory networks involving Psmd13, miR- 29a, and Dicer in various stages of neuronal differentiation and disease contexts.

## Materials availability

The requests for materials and reagents used in this protocol can be directed towards [D.K].

## Data and Code Availability

The ChIP sequencing data that support the findings of this study are openly available in Gene Expression Omnibus (GEO), reference number: GSE282821 (token code: azypagiiftuvnaj) and small RNA-seq datasets are available in GEO, reference number: GSE287514 (token code: obkxascmxlchrst).

All the softwares used in this study is listed in **Table S5**.

Upon a reasonable request, the corresponding authors of this article will provide unrestricted access to the original data" in the manuscript.

## Experimental Procedures

### Ethics Statement

All experimental protocols were approved by the Institutional Animal Care and Use Committee at the Monash University (Ethics ID: 22020).

All methods were carried out in accordance with relevant guidelines and regulations.

All methods are reported in accordance with ARRIVE guidelines.

### Cell lines

HEK293T (Human embryonic kidney) cells were kindly provided by Polo Lab. HEK293T cells were cultured in Dulbecco’s Modified Eagle Medium (DMEM) (Cat# 11995065, Gibco) supplemented with 10% fetal bovine serum, 2mM GlutaMAX Supplement (Cat# 35050061, Gibco), 100 I.U./mL Penicillin and 100 mg/mL Streptomycin.

### Collaborative Cross mouse model

Collaborative Cross mouse lines (CC) were obtained from Tel Aviv University (TAU), Israel and Geniad, Australia. Brain samples of CC strains were used in this study. We have used 22 CC lines from TAU and 32 lines from Geniad totalling to 54 CC lines. There were 3 CC mice under each strain. All the mice studied in this project were males. See **Table S1** for the list of CC mice and C57BL/6 strains used.

### Hippocampus dissection of CC brain

Frozen brain samples were removed using a long micro spatula to a petri dish containing RNAlater-ICE solution to protect the tissue from RNase. The brain was transferred to a wet filter paper and cut using a razor blade through the middle of the tissue. Hippocampus was dissected gently using two short spatulas. In brief, the spatula tips were positioned near the junction between the cortex and cerebellum, the cortical hemisphere was peeled off to expose the hippocampus. Carefully the extra white matter surrounding the hippocampus was removed. Holding the brain with one tip and placing the other tip just under the caudal side, hippocampus was rolled off from the remaining tissue.

### Mouse NPC culture

mNPCs were maintained on tissue culture-treated polystyrene plates with cell culture qualified Poly-L ornithine (PLO) (Cat# P4957, Sigma)/ Laminin (Cat# L2020, Sigma) solution in DMEM (Cat# 10565018, Gibco). The mNPCs were split every 3 – 4 days using Accutase (Cat# 07920, Stem Cell Technologies). The NPCs utilized for the experiments were between passages 2 and 8.

### Mouse primary neural progenitor cell isolation

Mouse Neural progenitor cells (mNPCs) were isolated from hippocampus of the WT C57BL/6 male mice aged between 30-40 weeks^28^. Briefly, animals were euthanized in a CO_2_ chamber. The skin around the head region was removed to expose the skull. It was cut open with small scissors without damaging the brain The brain was removed using a spatula and transferred to ice-cold HBSS-Hepes solution. The two hemispheres were separated, and the surrounding tissue was cleared until hippocampus was exposed. Hippocampus was carefully detached and stored in 5ml HBSS-Hepes solution. mNPCs were isolated by adding 5ml of Dissociation media to the hippocampus slices. This mixture was incubated at 37°C for 15 min. Next, the tissue was triturated 10 times with a 5mL plastic pipette to dislodge the pellet and was incubated again for 15 min at 37°C. Then 1 volume of ice-cold Solution 3 was added to inactivate the Trypsin and mixed gently by pipetting up and down using a 10mL plastic pipette. The cells were passed through a 70 µm strainer into a 50 mL falcon tube and centrifuged for 5 min at 1300 rpm at 4°C. The supernatant was removed, and the cells were resuspended in 10 mL ice cold Solution 2. They were centrifuged for 10 min at 2000 rpm at 4°C and resuspended in 2 mL ice cold Solution 3. The centrifugation was repeated for 7 min at 1500 rpm at 4°C. The cells were finally resuspended in 1 mL NPC growth medium (Penicillin/Streptomycin (100 units/ml), HEPES (8 mM), B27, FGF (10 ng/ml), EGF (10 ng/ml), DMEM: F12/Glutamax).

### ChIP-seq assay, library preparation, and sequencing

ChIP-seq was performed as described previously in^29^ with slight modifications. Briefly, mNPCs were trypsinized and washed once in PBS. Cells were resuspended in PBS and crosslinked in 1% formaldehyde solution (Cat# 252549, Sigma) for 8 min at room temperature. Then glycine was added at a final concentration of 0.125M to quench the reaction for 5 min. Cells were washed twice in PBS and suspended in SDS-ChIP buffer (20mM Tris-HCl, pH 8, 150mM NaCl, 2mM EDTA, 0.1% SDS, 1% Triton X-100 and protease inhibitor (Cat# C12010011, Diagenode)). Then, chromatin was sheared using a Diagenode Bioruptor Plus with high power mode for 40 cycles (sonication cycle: 30 sec ON, 30 sec OFF) until DNA was fragmented to 200–700bp. Sonicated chromatin was centrifuged at 4°C for 10 min and pre-cleared using Dynabeads Protein A/G beads for 1hr at 4°C with end-to-end rotation. Cleared supernatant was incubated with gene-specific primary antibody (Psmd13, Dicer) at 4°C overnight with end-to-end rotation. Protein A/G beads were added to the overnight incubated antibody-protein complex for 2 hrs at 4°C with end-to-end rotation to immunoprecipitate the chromatin. This complex was washed six times (5 min/wash at 4°C with rotation) in Low-salt buffer (twice, 50 mM HEPES pH 7.5, 150mM NaCl, 1mM EDTA, 1% Triton X-100, 0.1% sodium deoxycholate), High-salt buffer (once, 50 mM HEPES pH 7.5, 500mM NaCl, 1mM EDTA, 1% Triton X-100, 0.1% sodium deoxycholate), LiCl wash buffer (once, 10mMTris-HCl pH 8.0, 1mM EDTA, 0.5% sodium deoxycholate, 0.5% NP-40, 250mM LiCl) and TE buffer (twice, 10mMTris-HCl pH 8.0, 1mM EDTA pH 8). The chromatin was eluted and reverse-crosslinked in SDS-Elution buffer (1% SDS, 50mMTris-HCl pH 8.0, 10mM EDTA pH 8) at 65°C overnight. ChIP DNA was treated with 1µl RNase (10mg/ml) for 1hr and 3µl Proteinase K (20mg/ml) for 3hrs at 37°C. The purified ChIP DNA and Input DNA (reserved before adding antibody but reverse crosslinked and purified) was used to prepare ChIP-seq libraries using NEBNext Ultra II DNA Library Prep Kit for Illumina (Cat. E7645S, New England Biolabs) DNA samples were ligated to adaptor oligos for multiplex sequencing (Cat. E7335G, New England Biolabs). ChIP-seq was performed using Illumina Hiseq3000/4000 sequencing platform.

### Immunocytochemistry

Cells were treated in different experimental conditions for specified time. Then, the cells were fixed in 4% paraformaldehyde (Cat#: Sc-281692, Santa Cruz Biotechnology) for 10min at room temperature. The plates were briefly rinsed in PBS and permeabilised for 30min at room temperature using permeabilization buffer (PBS-0.1% Tween20, 5% goat serum). The cells were incubated with specific primary antibody ß III Tubulin at 4°C overnight. The following day cells were washed with PBS for 10min/3washes. We added secondary antibody (Alexa Fluor 546 anti-Rabbit) to the cells and incubated for 1hr at room temperature in dark setup. We repeated the washing step with PBS for 10min/3washes protected from light. The cells were counterstained by DAPI for 5min at room temperature in dark and imaged using confocal SP5 5 channel microscope.

### ChIP-seq analysis

To analyze ChIP-seq data, initial processing involved trimming raw data with the trim galore program to eliminate adapter sequences. The program’s --paired function was utilized to validate paired-end reads, and low-quality sequence reads were removed. Quality assessment of the sequencing data was conducted using FastQC. The Mm10 reference genome was employed for alignment with Bowtie v2.3.5, and the output SAM/BAM files were converted to sorted BAM files using Samtools, duplicates removed using filterdup. Enriched regions (peaks) were identified through peak-calling algorithms MACS3. MACS3 callpeak function (macs3 callpeak -t treatment file -c Input file -g mm -n ChIPpeaks --nomodel -p 0.01) assessed the significance of enriched binding regions over the input control. Reproducible peaks from all samples were then merged to create a union peak set. Additional processed ChIP-seq alignments and peak calls were downloaded from ENCODE from the following accessions ENCSR129DIK, ENCSR352NVU and ENCSR428OEK. Peaks were annotated with genomic features using ChIPseeker. We employed 2000 bp regions surrounding the TSS to create density plots that illustrate the ChIP-seq signal for Psmd13 and Dicer. Correlation matrices and heatmaps were generated using deeptools (computeMatrix, plotHeatmap, plotCorrelation). DNA sequence motifs within the peaks were identified using motif discovery tool MEME-ChIP. ChIP-seq signal was converted to bigwig format for visualization using deepTools bamCoverage v 3.3.1 with the following parameters: --bs 5 --smoothLength 105 -- normalizeUsing RPKM. Visualization of aligned reads and identified peaks was achieved through genome browsers such as Integrative Genomics Viewer (IGV) and UCSC Genome Browser. Lastly, GO enrichment analysis was conducted using ClusterProfiler. To look for differentially accessible regions a consensus peak set using samples in different experimental conditions was produced using the R package DiffBind.

### Small RNA sequencing

Sequence libraries were filtered for adaptor contamination using the Cutadapt (v3.3) software tool. This software cut the adaptor sequence from the sequencing reads and filtered reads whose length was greater than or equal to 15 bp for further analysis. Filtered reads were aligned to the mouse reference genome (mm10) using Bowtie (v.2.3.5). Reads that mapped with two or fewer mismatches to the reference sequence were retained for further analysis. We used the re-annotated miR list (MirGeneDB) to count the mapped reads using the Featurecounts (v1.4.6) module from the subread package. These counts were normalized to library size, and we performed a differential expression analysis using the edgeR bioconductor package (v.3.8.6). The Limma (v3.60.4) package was used to calculate the differential expression change.

### Proteasome activity assay

mNPCs were treated with the proteasome inhibitor MG132 for different time intervals. Cells were then lysed in PBS with 0.5% NP-40 at 4°C and centrifuged at 10,000 rpm for 10 minutes at 4°C. The supernatants were collected and analyzed using a Proteasome Activity Assay Kit (Cat# ab107921; Abcam), following the manufacturer’s instructions.

### Co-IP and immunoblotting

Co-immunoprecipitation was performed using Dynabeads Co-Immunoprecipitation Kit (Cat#: 14321D, Life Technologies) following the manufacturer’s protocol. Before preparing the cells for co-IP, we coupled 5ug of antibody (Psmd13, Dicer and Rb igg) to the epoxy beads overnight at 4°C. Beads were washed as per the kit’s instruction. mNPCs were grown overnight at 37°C. The cells were lysed in detergent lysis buffer and washed once in PBS. The whole cell lysate was coupled to the antibody-bead complex and incubated at 4°C on a roller for 30min. This mixture was washed and eluted. The purified protein was subjected to immunoblotting.

For immunoblotting (western blotting), protein samples were subjected to SDS-PAGE. The samples were mixed with 2x Laemmli buffer (Cat#: 1610737, Bio-rad) and boiled for 5min at 95°C. It was resolved on 4–20% Mini-PROTEAN TGX Precast Protein Gel (Cat#: 4561093, Bio-rad). The protein gels were transferred using the iBlot2 mini gel Transfer Stack system (Cat#: IB24002, Thermo Fisher Scientific) and iBlot2 gel transfer device (Cat#: IB21001, Thermo Fisher Scientific) for 7 min at 25V. Membranes were blocked with 3% BSA in TBS-0.1%Tween 20 for 1hr at room temperature. The primary antibodies (Psmd13, Dicer and B-actin) were added to the membrane and incubated overnight at 4°C with continuous agitation. The membranes were washed in TBST for 3 times with 10min mixing between each washes. An HRP-conjugated secondary antibody (Anti-Rabbit IgG–Peroxidase) was added to the membrane for 1hr at room temperature. 10 min wash was repeated for 3 times with constant agitation. We detected the bands using SuperSignal West Atto (Cat#: A38554, Thermo Fisher Scientific).

### RNA extraction and real-time PCR analysis

RNA was isolated using miR Vana miR isolation kit following the manufacturer’s protocol. Briefly, cells were grown in different experimental conditions and harvested using 0.05% Trypsin-edta. Cells were washed in PBS and pellet lysed in lysis buffer provided with the kit. Oligonucleotides used in this method is listed in **Table S2**. For miR isolation, miR enrichment homogenate solution was added. For cDNA synthesis, iScript cDNA synthesis kit was used. For Taqman assays, we used Taqman advanced miR cDNA synthesis kit (**Table S3**). The PCR reaction was performed in Roche LC480 (Roche) using Kapa fast Sybr green Master mix for mRNA and Taqman Fast Advance master mix for miR. The gene expression data was normalised to Gapdh (mRNA) and mmu-mir-16/mmu-mir-191 (miR).

### siRNA transfection on mNPCs

For siRNA knockdown experiments, mNPCs were detached from the cell culture plates using Accutase. Then, centrifuged at 200rpm for 5min at room temperature, supernatant was removed, and pellet dissolved in complete growth medium. Cells were then distributed into pre-coated 24 well plates to perform knockdown. Transfection was performed directly before the cells attach to the surface of the plate using lipofectamine RNAimax (Cat. 13778075, Invitrogen) following the manufacturer’s protocol for 48hrs. For few of the genes, we performed forward transfection wherein the cells were allowed to grow overnight and then transfected the next day in a similar manner to increase the efficiency of transfection. For mNPC differentiation experiments, we grew cells and transfected them for 48hrs. Then re-transfected the cells again and differentiated them for three days. We targeted for knockdown efficiency above 70% for all the siRNAs used in this study.

### Pathway enrichment analysis for candidate gene prioritisation

Functional enrichment analysis for Gene Ontology (GO) terms was conducted using various bioinformatics platforms, including Panther, DAVID, and EBI, with Mus musculus as the background species. Modules enriched within the GO Biological Process were identified. Gene Ontology terms were selected based on a significance threshold of p-value < 0.05, adjusted using the Benjamini-Hochberg correction.

### Lentiviral transduction

All the lentivirus particles were purchased from Vector builder. We produced the second-generation lentivirus by transient transfection of HEK293T cells with lentiviral particles (Psmd13 and miR-29a; Table S4), pPax2 (packaging plasmid) and pMD2.G (envelope plasmid) in Optimem (Cat#: 31985070, Thermo Fisher Scientific) using Lipofectamine LTX reagent (Cat#: 15338100, Thermo Fisher Scientific). The complex was harvested after 72hrs, filtered through 0.45μm Durapore Membrane Filter (Cat#: S2HVU01RE, Merck Millipore) and concentrated using Amicon Ultra-15 Centrifugal Filter Units (Cat#: UFC910096, Merck Millipore). The mNPCs were infected using the lentiviruses and stable plasmid expression was generated using Puromycin selection.

### Luciferase assay

mNPCs were plated in 24 well PLO-Laminin coated cell culture plates for luciferase assay. mNPCs were co-transfected with siPsmd13 and miR-29a promoter expressing lentiviral. A mutant miR-29a promoter expressing lentiviral particles were also used. A scrambled lentiviral negative control was used to measure the background reporter activity. Cells were lysed 48hrs after transfection. Dual Luciferase Assay System (Cat#: E1910, Promega) was used to measure the luciferase activity according to the manufacturer’s instructions. The Firefly luciferase activity was analysed relative to the Renilla luciferase activity in the same sample by using a multi-mode microplate reader (FLUOstar Omega, BMG Labtek).

### Proteasomal activity assay

Proteasomal activity was measured using the Proteasome Activity Assay Kit (Abcam, Cat#: ab107921), following the manufacturer’s protocol. In brief, 10^6 cells were harvested and washed with cold PBS. The cells were then resuspended in 0.5% NP-40 in PBS and homogenized by pipetting. The homogenate was centrifuged at 13,000 rpm for 10–15 minutes at 4°C, and the supernatant was collected. For the assay, 100 μL of standard dilutions were used for the standard wells, MG132 treated (for the indicated times) and control samples for the sample wells, and 10 μL of positive control (in duplicate). The volumes were adjusted with assay buffer, followed by the addition of either the proteasome inhibitor or assay buffer, and 1 μL of proteasome substrate. The plate was incubated at 37°C, protected from light. Fluorescence was measured at 350/440 nm, followed by a second reading after 30 minutes if the sample activity was low.

### QTL analysis

The framework for QTL analysis was implemented using R. HAPPY package is available as an R package called happy.hbrem. To run happy in R, we also installed g.data and multicore.

We downloaded the condensed genome library from: http://mtweb.cs.ucl.ac.uk/mus/www/preCC/CC-2018/LIFTOVER/

happy.preCC.R script was obtained from

http://mtweb.cs.ucl.ac.uk/mus/www/preCC/R.CD/happy.preCC.R which was used to set the environment for mapping in R. miR expression profiling data was used to create phenotypic file. Then, condensed genome library was loaded which scanned for the phenotypic files to create an association between the trait in question and genetic markers at a particular genomic locus. Genome wide significance for miR trait were computed at 95% threshold. We also used Gene miner software to perform the QTL analysis.

### Quantification and statistical analysis

We used GraphPad Prism software to perform the quantitative analysis for all the experimental conditions. The normality of variables was determined by QQ-plot. For comparison between two datasets, we used student’s T-test. To test between multiple experimental variables, we performed One-way ANOVA. All the p-values permuted to p<0.05 were considered significant. The level of significance in all the measurements were represented as ∗ p < 0.05, ∗∗ p < 0.005, ∗∗∗ p < 0.0005, ∗∗∗∗ p < 0.0001. Statistical parameters specifying the total number of measurements (n), standard error for precision (mean ± SD) and p-values are reported in the individual figure legends.

## Acknowledgments

Confocal imaging was performed at the Monash Microimaging facility at Monash University, Clayton campus. This work was financially supported by the program of “Genetic diversity in the collaborative cross model recapitulates miRNA mediated neurogenesis” (ID: 281589068), which was a donation of Apex Biotech Research Pty Ltd to Monash University for Z.X.

## Author contributions

Conceptualization, Z.X. and D.K.; Methodology, D.K. and Z.X.; Validation, Z.X. and D.K.; Formal Analysis, D.K., G.M. and Z.X.; Investigation, D.K. and Z.X.; Software, D.K. and G.M.; Validation, D.K., G.M. and Z.X; Visualization, D.K., G.M. and Z.X.; Resources, G.M., D.K. and Z.X.; Project administration, D.K.; Writing – Original Draft, D.K.; Writing – Review & Editing, D.K. Z.X. and G.M.; Funding Acquisition, Z.X.; Supervision, Z.X.

