## Supplemental information for "Psmd13, a proteosome regulatory subunit identified in miR-29a regulation during neurogenesis"

Figure S1

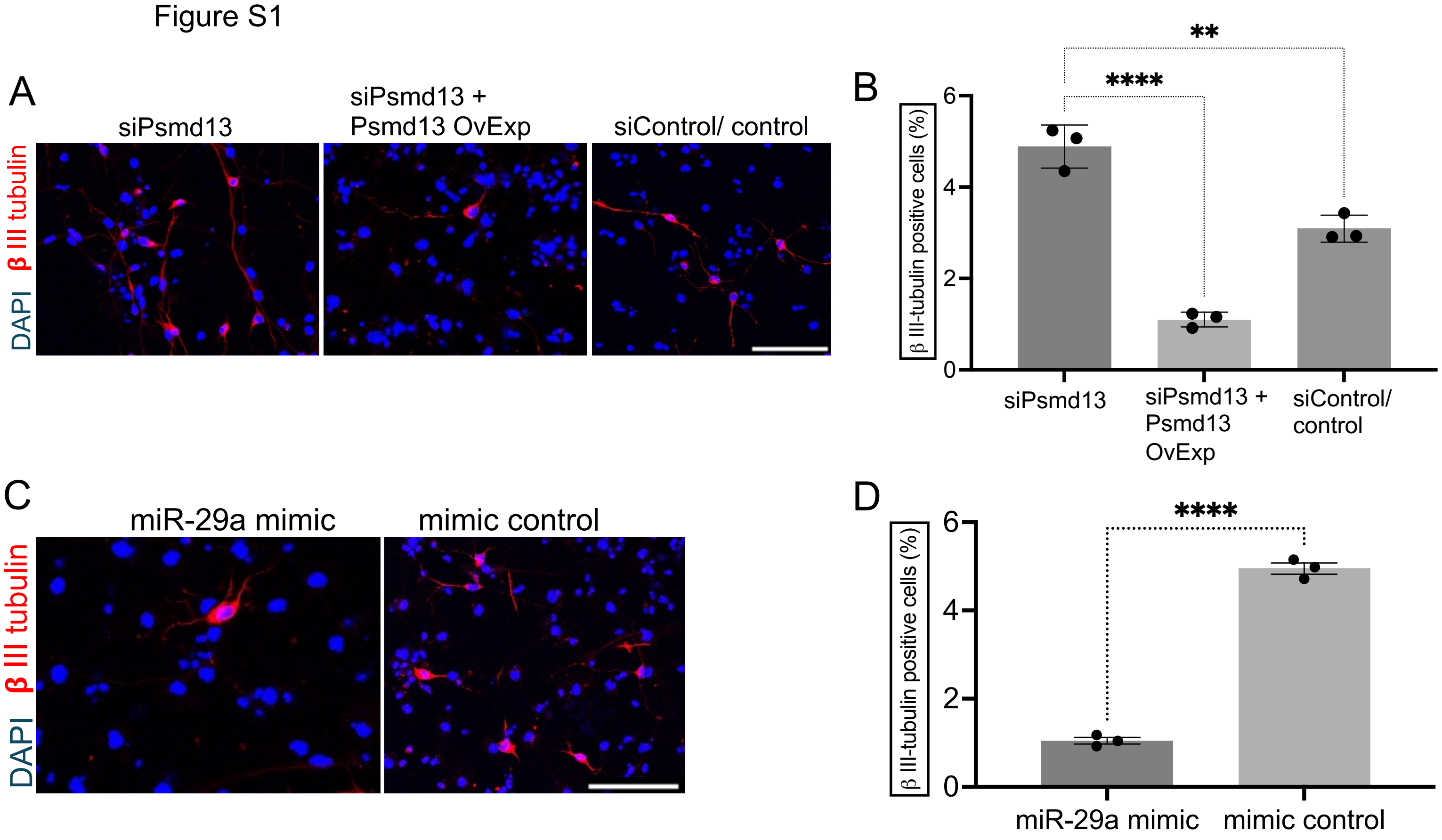
A
siPsmd13
siPsmd13 + Psmd13 OvExp
siControl/ control
DAPI β III tubulin
B
β III-tubulin positive cells (%)
6
4
2
0
****
**
siPsmd13
siPsmd13 + Psmd13 OvExp
siControl/ control
C
miR-29a mimic
mimic control
DAPI β III tubulin
D
β III-tubulin positive cells (%)
6
4
2
0
****
miR-29a mimic
mimic control

Figure S2

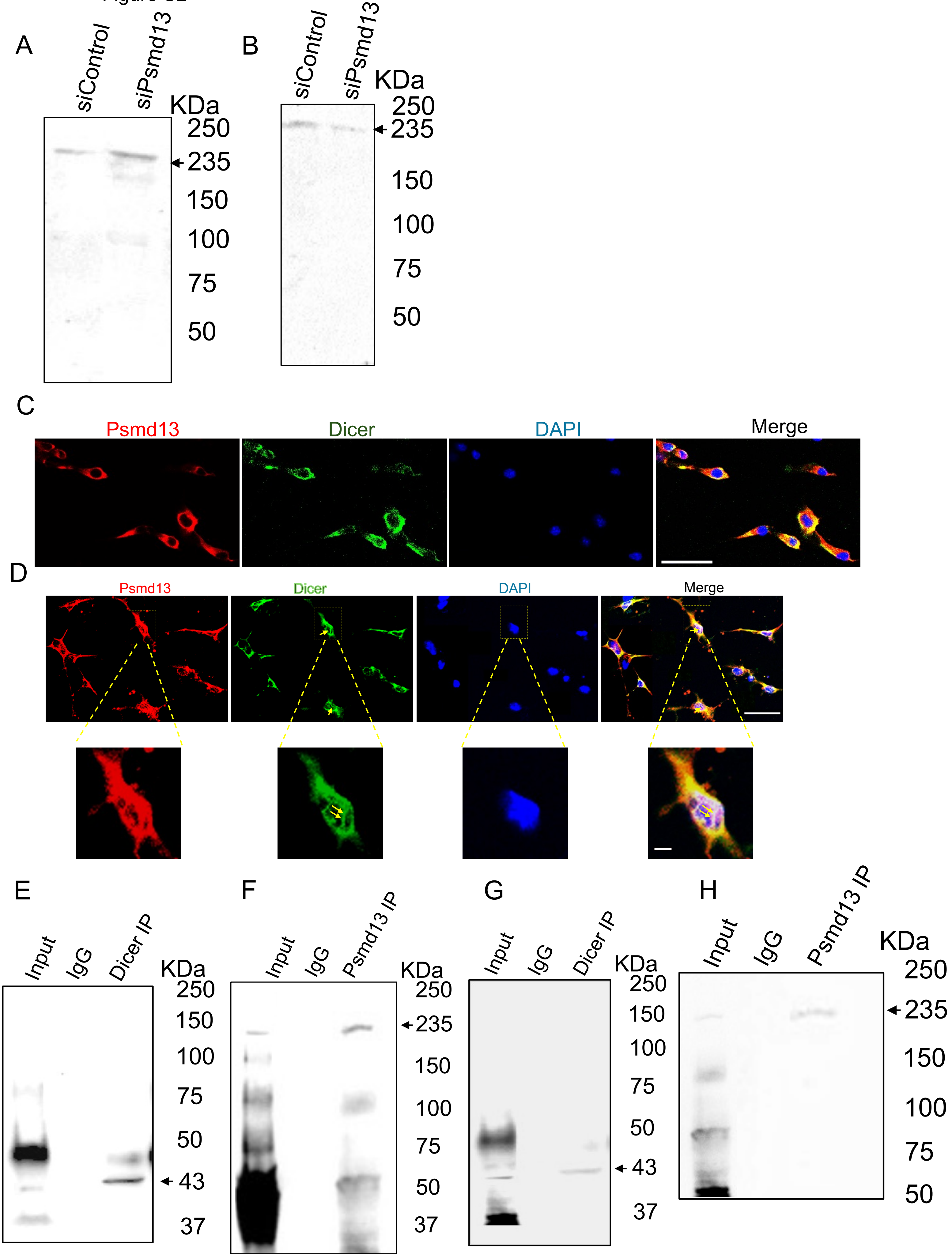
A
siControl
siPsmd13
KDa
250
235
150
100
75
50
B
siControl
siPsmd13
KDa
250
235
150
100
75
50
C
Psmd13
Dicer
DAPI
Merge
D
Psmd13
Dicer
DAPI
Merge
E
Input
IgG
Dicer IP
KDa
250
150
100
75
50
43
37
F
Input
IgG
Psmd13 IP
KDa
250
235
150
100
75
50
37
G
Input
IgG
Dicer IP
KDa
250
150
100
75
50
43
37
H
Input
IgG
Psmd13 IP
KDa
250
235
150
100
75
50

Figure S3

A

Proteins: 5800
Interactions: 52120

Undifferentiated

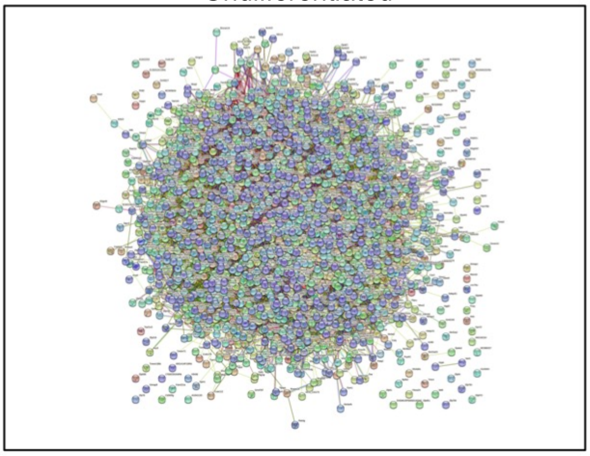

B

Proteins: 11615
Interactions: 92135

Differentiated

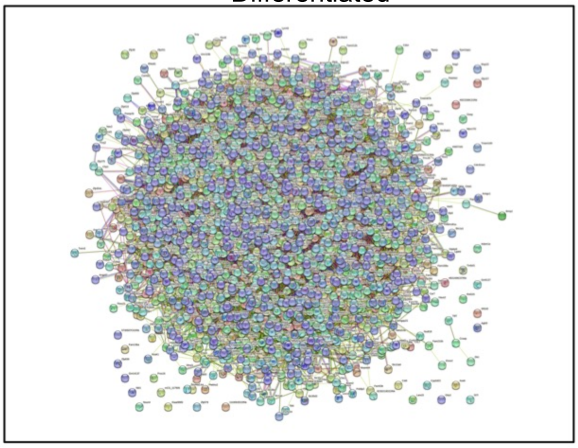

### Figure S4

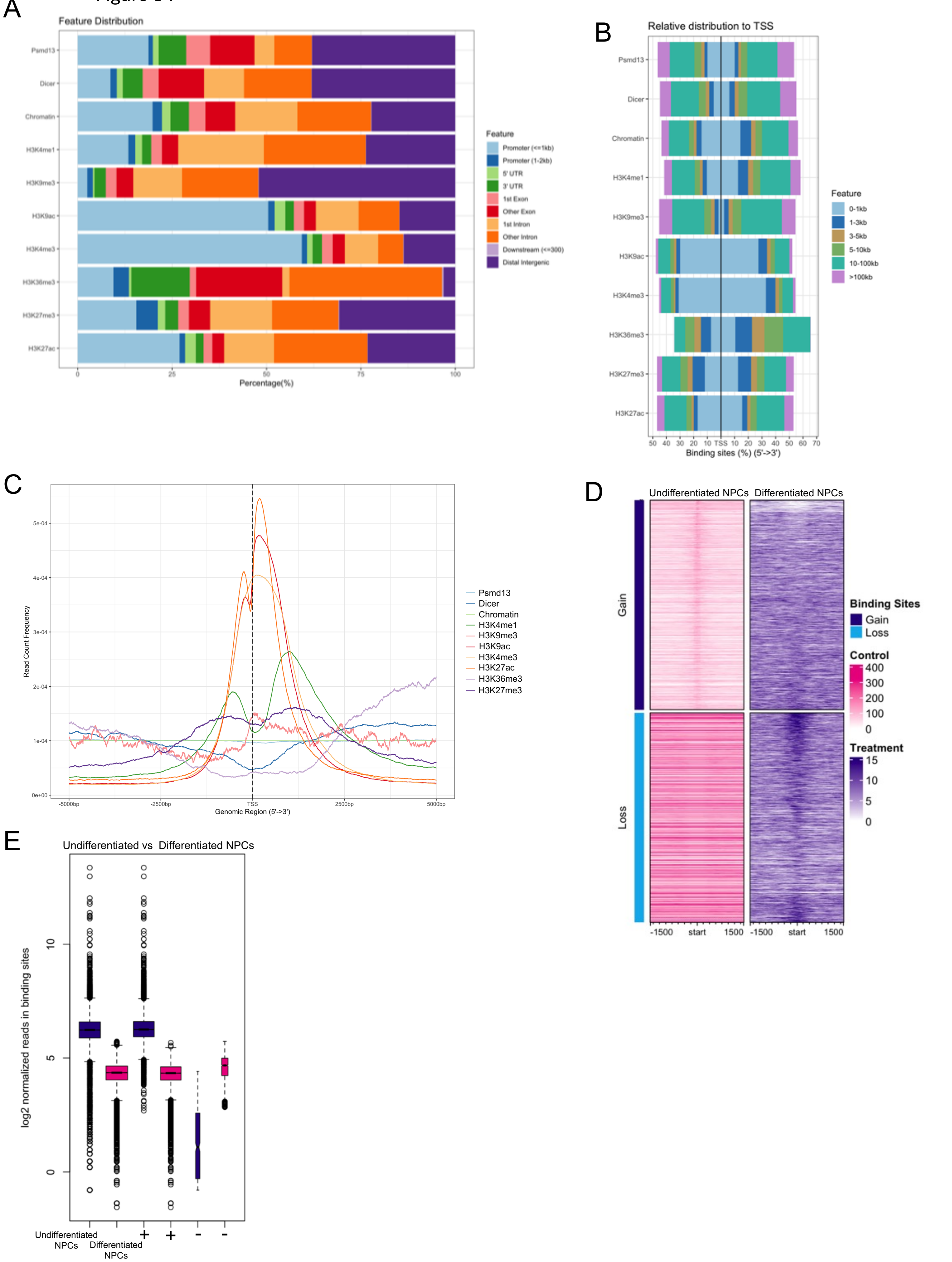


+ indicates sites with increased affinity in Undifferentiated NPCs
- indicates sites with increased affinity in Differentiated NPCs

Figure S5

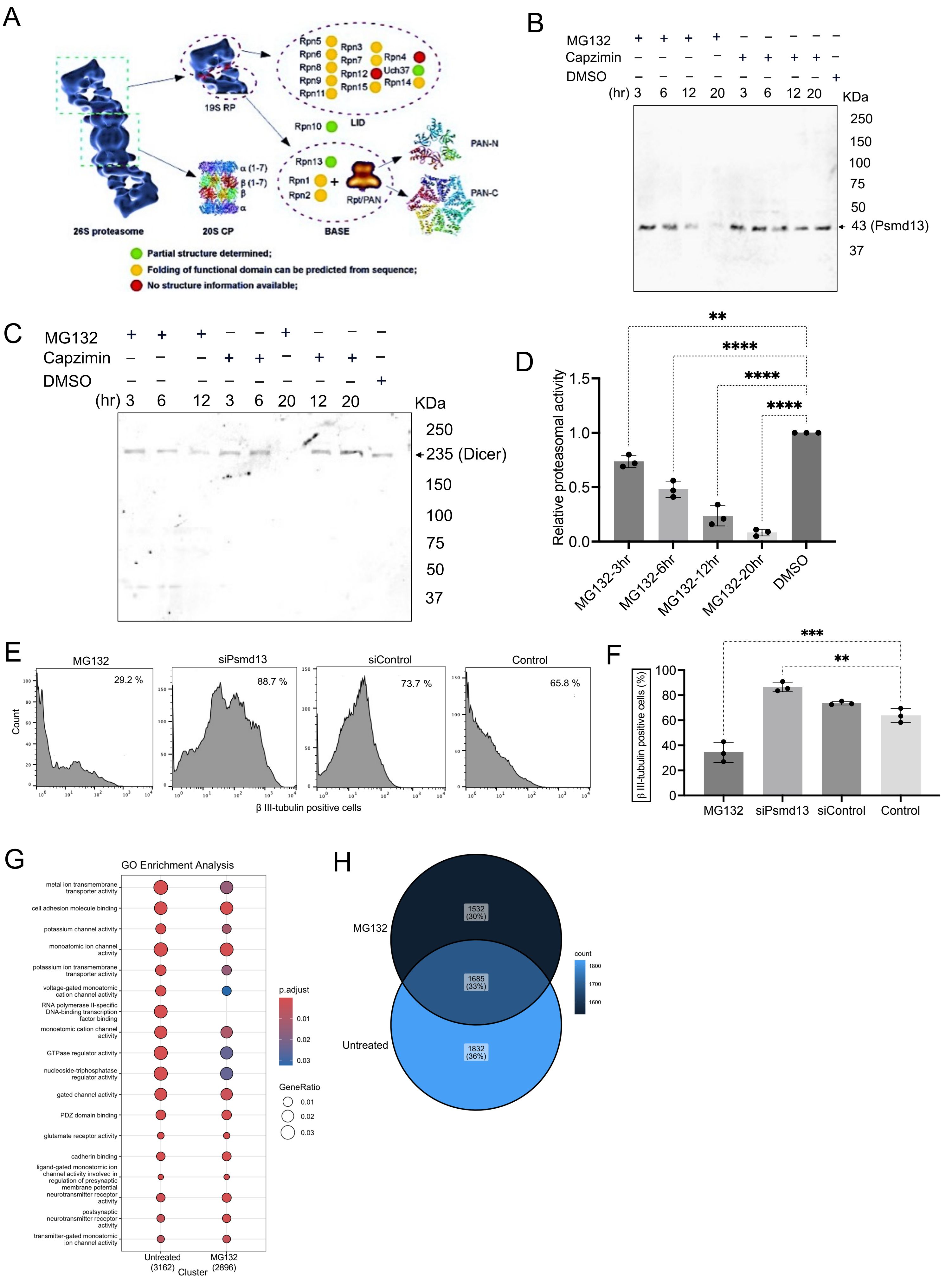
A
26S proteasome
19S RP
20S CP
Rpn5
Rpn6
Rpn8
Rpn9
Rpn11
Rpn3
Rpn7
Rpn12
Rpn15
Rpn4
Uch37
Rpn14
LID
Rpn10
Rpn13
Rpn1
Rpn2
Rpt/PAN
BASE
PAN-N
PAN-C
α (1-7)
β (1-7)
β
α
Partial structure determined;
Folding of functional domain can be predicted from sequence;
No structure information available;
B
MG132
Capzimin
DMSO
(hr) 3 6 12 20 3 6 12 20
KDa
250
150
100
75
50
43 (Psmd13)
37
C
MG132
Capzimin
DMSO
(hr) 3 6 12 3 6 20 12 20
KDa
250
235 (Dicer)
150
100
75
50
37
D
Relative proteasomal activity
1.5
1.0
0.5
0.0
MG132-3hr
MG132-6hr
MG132-12hr
MG132-20hr
DMSO
**
****
****
****
E
MG132
29.2 %
siPsmd13
88.7 %
siControl
73.7 %
Control
65.8 %
Count
β III-tubulin positive cells
F
β III-tubulin positive cells (%)
100
80
60
40
20
0
MG132
siPsmd13
siControl
Control
***
**
G
GO Enrichment Analysis
metal ion transmembrane transporter activity
cell adhesion molecule binding
potassium channel activity
monoatomic ion channel activity
potassium ion transmembrane transporter activity
voltage-gated monoatomic cation channel activity
RNA polymerase II-specific DNA-binding transcription factor binding
monoatomic cation channel activity
GTPase regulator activity
nucleoside-triphosphatase regulator activity
gated channel activity
PDZ domain binding
glutamate receptor activity
cadherin binding
ligand-gated monoatomic ion channel activity involved in regulation of presynaptic membrane potential
neurotransmitter receptor activity
postsynaptic neurotransmitter receptor activity
transmitter-gated monoatomic ion channel activity
Untreated (3162)
MG132 (2896)
Cluster
p.adjust
0.01
0.02
0.03
GeneRatio
0.01
0.02
0.03
H
MG132
Untreated
1532 (30%)
1685 (33%)
1832 (36%)
count
1800
1700
1600

**Figure S1**. Screening of upstream candidate genes using neuronal differentiation assay in mNPCs. Related to Figure 2.

(A-B) Immunostaining of differentiated mNPCs with Psmd13-depleted, Psmd13-depleted plus mNPCs overexpressing Psmd13 and control. Representative images (A) and quantification (B) of βIII-tubulin positive mNPCs. N = 3 experiments. mean ± SD, **p<0.01, ****p<0.0001, One-way Anova, Image scale bars=10 μm.
(C-D) Immunostaining of differentiated mNPCs with miR-29a mimics and mimic control. Representative images (C) and quantification (D) of βIII-tubulin positive mNPCs. N = 3 experiments. mean ± SD, ****p<0.0001, Unpaired T-test, Image scale bars=10 μm.

**Figure S2**. Psmd13 associates with Dicer and regulate miR-29a expression in mNPCs. Related to Figure 3.

(A) Full blots showing Dicer protein in extracts from control and Psmd13-depleted mNPCs under undifferentiated conditions. B-actin was used as a loading control.
(B) Full blots showing Dicer protein in extracts from control and Psmd13-depleted mNPCs under differentiated conditions. B-actin was used as a loading control.
(C) Confocal microscopy images showing the localization of Psmd13 (red) and Dicer (green) in the cytoplasm of mNPCs in the undifferentiated mNPCs. DNA was stained with DAPI (blue). Scale bar, 10 μm.
(D) Confocal microscopy images showing cytoplasmic and nuclear localization of Psmd13 (red) and Dicer (green, highlighted in arrows for nuclear stain) in the differentiated mNPCs. DNA was stained with DAPI (blue). Scale bar, 10 μm and 40 μm (Insert).
(E) Full blots showing Dicer Co-IP to detect endogenous Psmd13 proteins in undifferentiated mNPCs.
(F) Full blots showing Psmd13 Co-IP to detect endogenous Dicer proteins in undifferentiated mNPCs.
(G) Full blots showing Dicer Co-IP to detect endogenous Psmd13 proteins in differentiated mNPCs.
(H) Full blots showing Psmd13 Co-IP to detect endogenous Dicer proteins in differentiated mNPCs.
Input and IgG antibody was used as controls for the experiment.

**Figure S3.** Psmd13 depletion impacts global miR regulation in mNPCs. Related to Figure 4.
(A) Integrated network of miRs and their predicted gene targets which are differentially expressed in the undifferentiated mNPCs. miR–mRNA interaction network constructed using Multimir containing DEMs and their validated miR gene targets.
(B) Integrated network of miRs and their predicted gene targets which are differentially expressed in the differentiated mNPCs. miR–mRNA interaction network constructed using Multimir containing DEMs and their validated miR gene targets.

**Figure S4**. Psmd13 dependency for Dicer binding at miR-29a locus in mNPCs. Related to Figure 5.

(A) Bar plot of the relative distribution of Psmd13, Dicer and Histone mark peaks to TSS.
(B) Bar plot of the percentage of annotated features of Psmd13, Dicer and Histone mark peaks.
(C) Metaplot of Psmd13, Dicer and Histone mark peaks around TSS.
(D) Heatmaps showing the enrichment of Dicer reads between undifferentiated and differentiated mNPCs in a 1500 bp window centered around the TSS. Scale is as indicated in the signal.

(E) Box plots of read distributions for significantly differentially bound sites in the undifferentiated and differentiated mNPCs.

**Figure S5**. Impact of proteasome inhibition on Dicer levels and miR-29a Expression in mNPCs. Related to Figure 6.

(A) Schematic representation of the 26S proteasome.
(B) Full blots showing the levels of Psmd13 protein after treatment with 5 μM MG132, 0.2 μM Capzimin and DMSO control in mNPCs for the indicated time duration
(C) Full blots showing the levels of Dicer protein after treatment with 5 μM MG132, 0.2 μM Capzimin and DMSO control in mNPCs for the indicated time duration.
(D) Quantification of proteasomal activity in mNPCs after treatment with MG132 for the indicated time duration. N = 3 experiments, mean ± SD, **p<0.01, ****p<0.0001. One-way Anova.
(E-F) Representative histograms (E) and quantification (F) of βIII-tubulin detection using flow cytometry in MG132 treated, untreated and non-targeting control differentiated mNPCs. N= 3 experiments. mean ± SD, **p<0.01, ***p<0.001. One-way Anova.
(G) The top 20 enriched GO terms of molecular function. Circle size indicates the number of genes enriched in each term. Color saturation represents the significance level.
(H) Overlap of the genes annotated between MG132 and untreated ChIP-seq profile displayed as Venn diagram. The P-value was calculated using Fisher's exact test.
